# A sequential, RNA-derived, modified adenosine pathway safeguards cellular metabolism

**DOI:** 10.1101/2024.05.23.595515

**Authors:** Akiko Ogawa, Satoshi Watanabe, Tomoyoshi Kawakami, Allen Yi-Lun Tsai, Jirio Fuse, Kimi Araki, Shinichiro Sawa, Kenji Inaba, Fan-Yan Wei

**Author notes:** Correspondence (A.O.) & (F.Y.-W.). These authors contributed equally (Satoshi Watanabe, Tomoyoshi Kawakami).

## Abstract

RNA contains diverse post-transcriptional modifications and its catabolic breakdown yields numerous modified nucleosides that must be properly processed, but the molecular mechanism is largely unknown. Here, we show that three RNA-derived modified adenosines, *N*^6^-methyladenosine (m^6^A), *N*^6^,*N*^6^-dimethyladenosine (m^6,6^A), and *N*^6^-isopentenyladenosine (i^6^A), are sequentially metabolized to inosine monophosphate (IMP) to prevent their intrinsic cytotoxicity. These modified adenosines are phosphorylated by adenosine kinase (ADK) followed by adenosine deaminase-like (ADAL)-mediated deamination in both plants and animals. ADAL knockout mice accumulate *N*^6^-modified AMPs that allosterically inhibit AMP-activated protein kinase (AMPK), leading to dysregulation of glucose metabolism. Furthermore, ADK deficiency, reported in patients with severe metabolic defects, induces aberrant elevation of m^6^A/m^6,6^A/i^6^A, disrupting lipid metabolism and causing early death in mouse models. The findings unveil a fundamental mechanism by which cells alleviate the toxicity of modified adenosines, and that connects modified adenosines to human disease.

**In brief:** RNA catabolism yields diverse nucleosides with modifications attached. Three RNA-derived modified adenosines (m^6^A, m^6,6^A, and i^6^A) are intrinsically toxic and subjected to sequential metabolism to yield inosine monophosphate. Dysregulation of this pathway impairs energy balance and contributes to metabolic diseases.

**Highlights:**

- Cytotoxic RNA catabolism-derived modified adenosines undergo sequential metabolism
- m^6^A, m^6,6^A, and i^6^A are phosphorylated by ADK then deaminated by ADAL to IMP
- m^6^AMP, m^6,6^AMP, and i^6^AMP allosterically inhibit AMP-mediated AMPK activation
- Loss of ADK causes the accumulation of m^6^A, m^6,6^A, and i^6^A, and abnormal lipid metabolism, which can lead to genetic disease

## INTRODUCTION

RNA modifications, collectively referred to as the epitranscriptome, control post-transcriptional gene expression and govern fundamental biological phenomena^1^, ^2^ including embryonic development^3^, immune responses^4^, hormone secretion^5^, tumorigenesis^6^, and learning and memory^7^. Aberrant modifications can profoundly affect human health and cause disease^8^. More than 170 types of RNA modifications have been identified in all three domains of life^9^. They play an essential role in post-transcriptional regulation of gene expression by controlling intracellular RNA localization^10^, RNA structure^11^, decoding capacity^12^, alternative splicing^13^, and stability^14^.

RNA molecules (RNAs) are catabolized to nucleotides or nucleosides as a consequence of turnover. The resulting unmodified nucleotides or nucleosides are subject to the salvage pathway for nucleotide recycling, or the degradation pathway to uric acid and β-alanine^15^. The purine/pyrimidine salvage pathway has been extensively studied for decades and is well established^16^. For example, adenosine, one of the four building blocks of RNA, is phosphorylated by adenosine kinase (ADK) to adenosine monophosphate (AMP), and subsequently converted to adenosine triosephosphate (ATP). Adenosine can also be deaminated by adenosine deaminase (ADA) to inosine, which is converted to inosine monophosphate (IMP) through a series of enzymatic reactions. IMP is then converted to guanosine monophosphate (GMP) or AMP, which can be used for the regeneration of GTP or ATP. Similarly, uridine and cytidine are also subject to a phosphorylation-dependent salvage pathway to generate cytidine triphosphate (CTP) or uridine triphosphate (UTP). Because salvage pathway-derived nucleotides constitute a large fraction of the intracellular nucleotide pool, many genetic diseases manifesting as neurological or metabolic disorders have been reported in individuals with pathogenic mutations in genes related to nucleotide metabolism.

In addition to unmodified nucleosides, RNA catabolism produces numerous nucleosides with modifications attached to the base or ribose groups^17,18,19^ whose biological roles are largely unknown. We recently revealed that these modified nucleosides are released from cells, partially through equilibrative nucleoside transporters (ENT1 and ENT2)^20^. Accumulation of modified nucleosides in ENT1/2-deficient cells induces an autophagic response, demonstrating that excessive accumulation of intracellular modified nucleosides is generally unfavorable. Interestingly, some modified nucleosides function as signaling molecules in the extracellular environment. For example, *N*^6^-methyladenosine (m^6^A), which is mainly derived from lysosomal degradation of ribosomal RNA (rRNA) and messenger RNA (mRNA), can activate adenosine A3 receptor with high potency and efficacy, and exaggerate inflammatory responses in mice^21^. Although normally maintained at low levels^21^, m^6^A can be rapidly upregulated upon cytotoxic stimulation. Thus, levels of intracellular and extracellular RNA-derived modified nucleosides are both dynamically and tightly regulated, and aberrant elevation can trigger various cellular responses, potentially leading to pathological consequences. However, the biological significance and regulatory mechanisms of most RNA-derived modified nucleosides remain elusive.

In this study, we performed functional and metabolic screening of RNA-derived modified nucleosides and elucidated an evolutionarily conserved mechanism specific for the three types of modified adenosines m^6^A, *N*^6^,*N*^6^-dimethyladenosine (m^6,6^A), and *N*^6^-isopentenyladenosine (i^6^A). These are the most intrinsically toxic among RNA-derived modified nucleosides, and are dynamically metabolized in cells and *in vivo*. We show that m^6^A, m^6,6^A, and i^6^A are phosphorylated by ADK to monophosphate forms, followed by deamination to IMP by adenosine deaminase-like (ADAL) in both animals and plants. ADAL deficiency resulted in accumulation of monophosphate forms of modified adenosines, which allosterically inhibit the activity of AMP-activated protein kinase (AMPK), leading to dysregulation of carbohydrate metabolism. Moreover, ADK deficiency resulted in the accumulation of modified adenosines, leading to defective lipid metabolism and early mortality in mice. Importantly, human disease-related ADK mutants displayed elevated m^6^A, m^6,6^A, and i^6^A, which potentially explains the catastrophic pathogenesis. Taken together, our findings reveal an evolutionarily conserved defense mechanism against the intrinsic toxicity of the three modified adenosines derived from RNA catabolism, and link the modified adenosines to genetic diseases.

## RESULTS

### m^6^A, m^6,6^A, and i^6^A possess intrinsic cytotoxicity and are rapidly metabolized *in vivo*

To comprehensively understand the biological roles of modified nucleosides, we treated multiple human cell lines including U87-MG (U87), MCF-7, MDA231, HEK293A, HepG2, and THP-1, and human primary cells (human conjunctival fibroblasts: HConF) with 43 species of modified nucleosides. Of these, 37 are modified nucleosides present in eukaryotes, five are modifications found in non-eukarya domain organisms, and two are secondary purine metabolites^22^. While most modified nucleosides had no effect or even slightly promoted cell viability, m^6,6^A, i^6^A, and m^6^A significantly reduced cell viability and exhibited cytotoxicity in a dose-dependent manner (Fig. 1A−C, Fig. S1A−H). Notably, i^6^A and m^6,6^A showed the strongest toxicity in all cell types tested in this study, while the toxicity of m^6^A varied with cell type (Fig. S1A−G).

**Figure 1.**
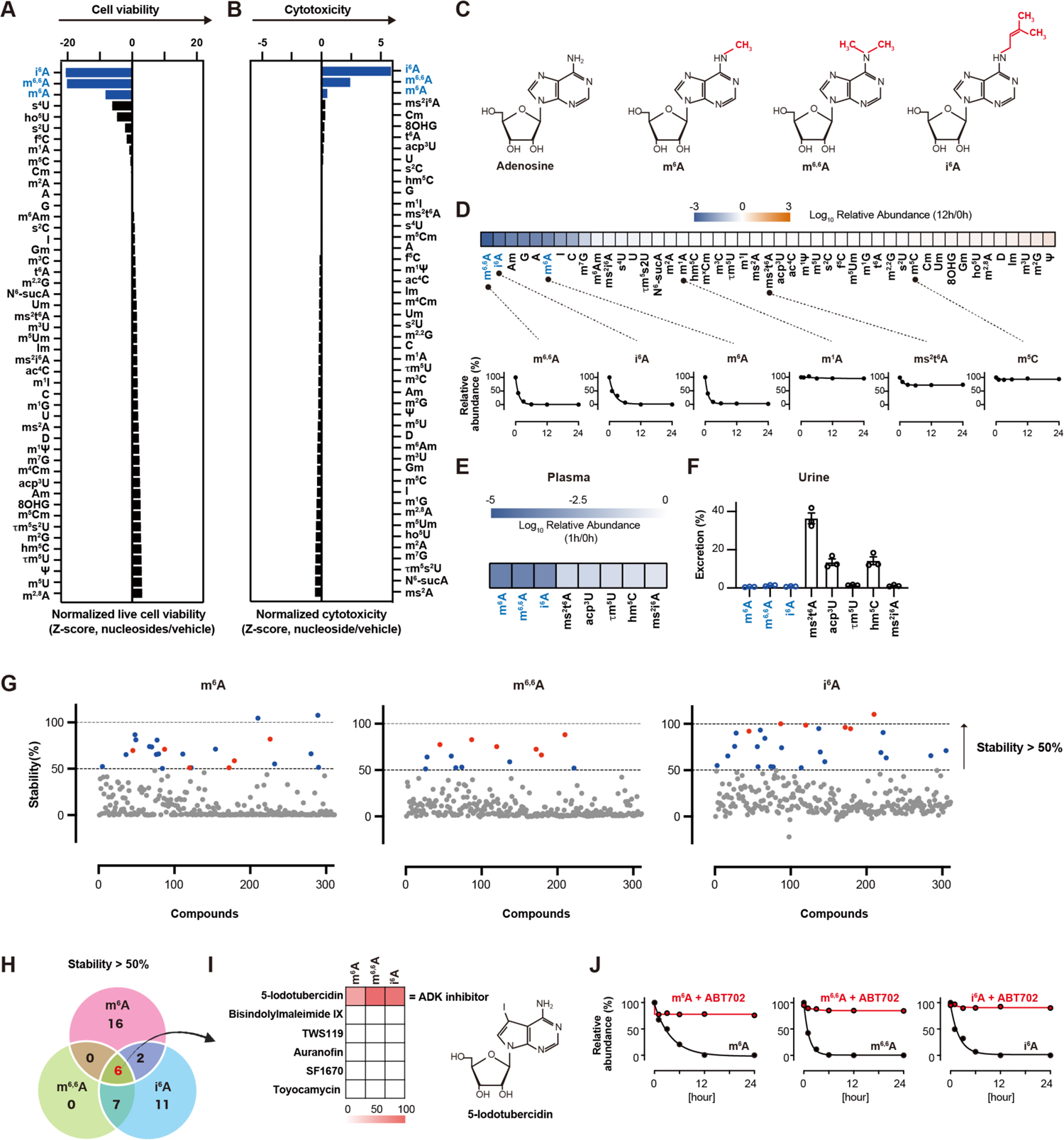
*N*^6^-methyladenosine (m^6^A), *N*^6^,*N*^6^-dimethyladenosine (m^6,6^A), and *N*^6^-isopentenyladenosine (i^6^A) possess intrinsic cytotoxicity and are rapidly metabolized. (A) Normalized cell viability of multiple human cell lines (U87, MCF7, MDA231, HEK293A, HepG2, THP1, and HConF) after 24 h incubation with each modified nucleoside (500 µM). The sum of the results from the seven cell lines is shown. The results for each cell are shown in Figures S1A−G. n = 3−16. Abbreviations for modified nucleosides are listed in Table S1. (B) Normalized cytotoxicity of HConF after 24 h incubation with each modified nucleoside (100 µM). n = 3−6. (C) Chemical structures of m^6^A, m^6,6^A, and i^6^A. (D) (upper panel) Heatmap showing the relative clearance of exogenously added modified nucleosides (1 µM) from the supernatants of HEK293 cells after 12 h compared to 0 h. Results are averages of n = 3. (lower panel) Time course of clearance of exogenous nucleosides from HEK293A supernatants (n = 3). Results are means ± standard error of the mean (SEM). (E, F) Mice were injected with a modified nucleoside mixture to achieve a final plasma concentration of 5 µM, and plasma and urine were collected 1 h after injection (n = 3). (E) Heatmap showing the plasma clearance of exogenously injected modified nucleosides. (F) Urinary excretion rate of modified nucleosides (% of total injection abundance). (G) Compound library screen in HEK293A cells. m^6^A, m^6,6^A, and i^6^A were added to HEK293A cells (1 µM) after 1 h treatment with 311 compounds (10 µM), and the supernatant was collected after 12 h to determine the stability of m^6^A, m^6,6^A, and i^6^A. Compounds that stabilized m^6^A, m^6,6^A, and i^6^A by >50% are indicated by colored dots, with red dots indicating those common to all three modified adenosines and blue dots for others. (H) Compounds stabilizing all three modified nucleosides (m^6^A, m^6,6^A, and i^6^A). (I) Secondary screening using six compounds at a lower concentration of 100 nM. (J) Time course of changes in the relative abundance of exogenously added m^6^A, m^6,6^A, and i^6^A (1 µM) in the supernatant of HEK293A cells in the presence or absence of ABT702 (selective adenosine kinase inhibitor, 10 µM). n = 3, results are means ± SEM.

Under normal cell culture conditions, extracellular levels of m^6^A, m^6,6^A, and i^6^A were low compared to their abundance in RNA, but they could be rapidly elevated upon cytotoxic stimulation (Fig. S1I−K). Intriguingly, when we applied modified nucleosides to cells, extracellular m^6^A, m^6,6^A, and i^6^A decreased by nearly 90% within 24 h, while other modified nucleosides showed much slower decrease (Fig. 1D, S1L).

Next, we intravenously administrated modified nucleosides including m^6^A, m^6,6^A, and i^6^A into mice and examined the turnover *in vivo*. Consistent with *in vitro* experiments, exogenous m^6^A, m^6,6^A, and i^6^A were rapidly cleared from mice plasma compared with other modified nucleosides (Fig. 1E). Importantly, the rapid disappearance of m^6^A, m^6,6^A, and i^6^A cannot be explained by urinary excretion, because their abundances were not elevated in urine when their serum levels declined (Fig. 1F). It should be noted that extracellular Am and m^7^G also rapidly disappeared after exogenous addition, although they did not possess any cytotoxicity (Fig. 1A-B). Taken together, these results show that m^6^A, m^6,6^A, and i^6^A possess intrinsic cytotoxicity and are subject to selective metabolic regulation to maintain their steady state at low levels in mammalian cells.

### ADK is responsible for metabolizing m^6^A, m^6,6^A, and i^6^A

To understand the metabolic regulation of m^6^A, m^6,6^A, and i^6^A, we performed unbiased screening by applying a metabolic enzyme inhibitor library to HEK293 cells in combination with exogenous m^6^A, m^6,6^A, and i^6^A. For the first screening, we applied 311 compounds at a fixed concentration of 10 µM to cells, followed by addition of m^6^A, m^6,6^A, and i^6^A. We identified six compounds that commonly suppressed the decrease in m^6^A, m^6,6^A, and i^6^A more than 2-fold compared with vehicle (Fig. 1G-H). For the secondary screening, we applied the compounds to cells at a lower concentration (100 nM). Among the six compounds, 5-iodotubercidin, an ADK inhibitor, almost completely blocked the decrease in exogenous m^6^A, m^6,6^A, and i^6^A (Fig. 1I). Blockade of m^6^A, m^6,6^A, and i^6^A metabolism was further confirmed by ABT702, another selective ADK inhibitor (Fig. 1J).

In mammalian cells, adenosine is subject to a complex metabolic pathway including the methionine cycle and folate cycle-mediated adenosine generation^23–24^, ADA-mediated deamination, and ADK-mediated phosphorylation (Fig. S2A). To investigate the role of ADK in the context of adenosine metabolism, inhibitors of folate, ADK, and ADA were applied to cells in the presence of various modified nucleosides. In line with screening results, the ADK inhibitor, but not ADA or folate inhibitors, attenuated the decrease in m^6^A, m^6,6^A, and i^6^A (Fig. S2B). Unexpectedly, we found that ADA inhibition suppressed the decrease in Am, and that neither ADA, ADK, nor folate inhibition rescued m^7^G levels, suggesting that Am is metabolized in a deamination-dependent manner and m^7^G by an unknown mechanism. (Fig. S2B).

Next, we generated ADK knockout (KO) cells (Fig. S2C−E) and analyzed m^6^A, m^6,6^A, and i^6^A levels. In ADK KO cells, steady-state levels of m^6^A, m^6,6^A, and i^6^A were highly elevated in the extracellular medium compared with parental wild-type (WT) cells, and adenosine was moderately elevated in the KO cells, while other modified nucleosides showed no difference between WT and KO cells (Fig. 2A-B). Two isoforms of ADK, different at the N-terminus, have been identified in mammals (Fig. S2F), and both are active with comparable biochemical or kinetic properties^25, 26^. Reintroduction of either ADK isoform significantly suppressed the accumulation of endogenous (Fig. 2B) and exogenously added (Fig. S2G) m^6^A, m^6,6^A, and i^6^A in ADK KO cells. Consistent with extracellular m^6^A, m^6,6^A, and i^6^A, their intracellular levels were also elevated in ADK KO cells, and were effectively suppressed in KO cells in which ADKs were reintroduced (Fig. S2H). It should be noted that in contrast to the modified adenosines, levels of unmodified adenosines were only moderately elevated in ADK KO cells both extracellularly and intracellularly (Fig. 2B, S2H).

**Figure 2.**
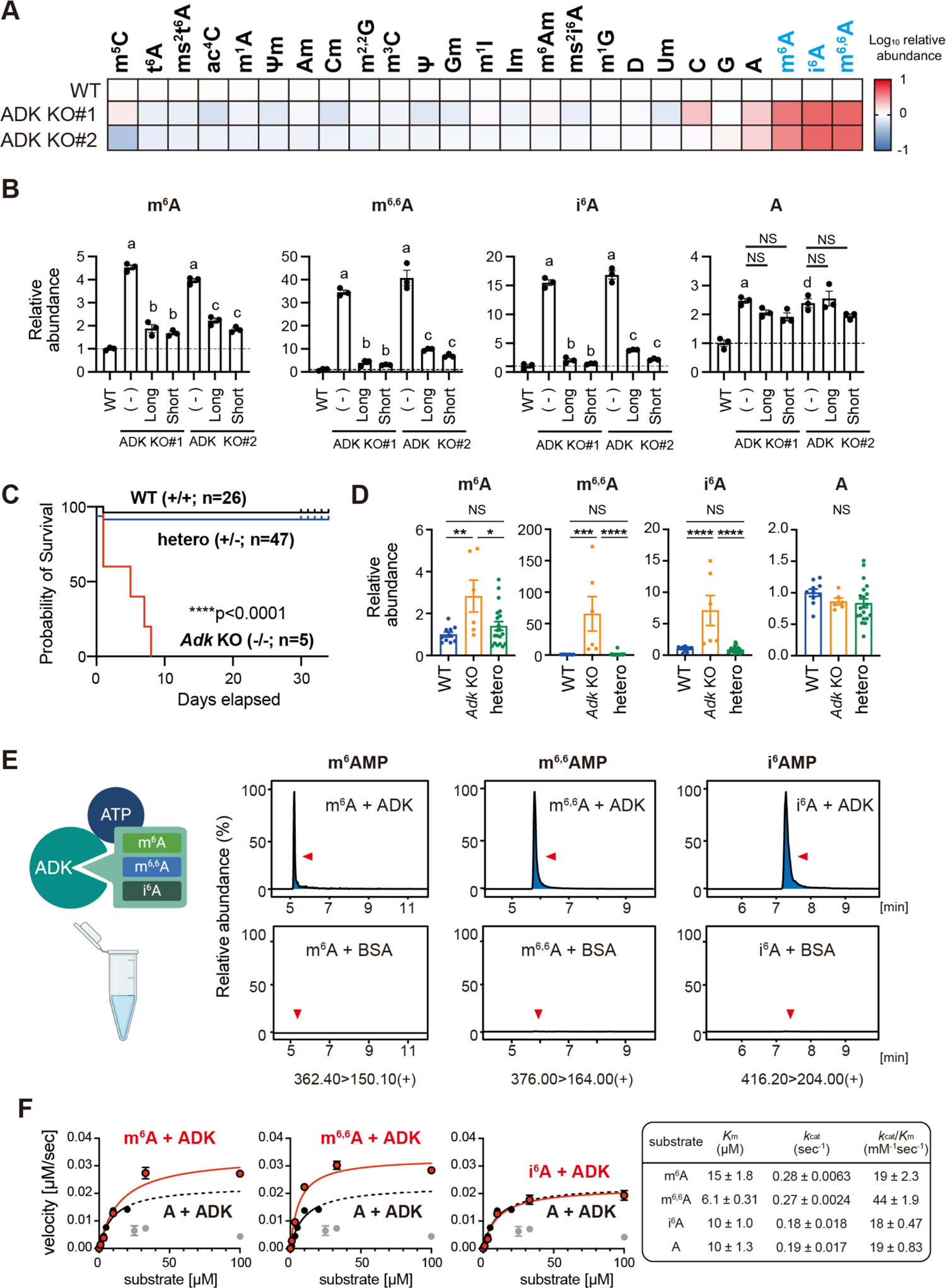
m^6^A, m^6,6^A, and i^6^A are substrates for adenosine kinase (ADK). (A) Heatmap showing the relative abundance of nucleosides in the supernatant of two clones of ADK KO cells compared to WT. Results are averages of n = 3. (B) Relative abundance of m^6^A, m^6,6^A, i^6^A, and unmodified adenosine in the supernatants of ADK KO cells and ADK KO cells following reintroduction of exogenous ADK (compared to the amount in the supernatant of WT cells). a, *p* <0.0001 vs. WT; b, *p* <0.0001 vs. ADK KO#1; c, *p* <0.0001 vs. ADK KO#2; d, *p* <0.001 vs. WT, by one-way analysis of variance (ANOVA) followed by Tukey’s multiple comparison test. (C) Kaplan-Meier survival curve of WT (black), heterozygous (blue), and *AdK^-/-^* (KO; red) mice. The overall log-rank *p* <0.0001 is when all three groups were calculated simultaneously. (D) Endogenous levels of m^6^A, m^6,6^A, and i^6^A in the liver of WT and *AdK* KO mice. \**p* <0.05 by unpaired t-test; \*\*\*\**p* <0.0001; \*\*\**p* <0.001; \*\**p* <0.01; and \**p* <0.05 by one-way ANOVA followed by Tukey’s multiple comparison test. (E) Relative abundance of m^6^AMP, m^6,6^AMP, and i^6^AMP measured by liquid chromatography-mass spectrometry (LC-MS) following *in vitro* phosphorylation assay using human ADK protein and control (bovine serum albumin: BSA). (F) Michaelis-Menten kinetic curves (left) and kinetic constants (right) of ADK for adenosine, m^6^A, m^6,6^A, and i^6^A (n = 3). Substrate inhibition occurred for adenosine at concentrations higher than 20 µM (plotted in gray), which were not used for calculation. Results are means ± SEM.

We further generated *Adk* KO mice to investigate the role of ADK in m^6^A, m^6,6^A, and i^6^A metabolism *in vivo* (Fig. S2I, S2J). *Adk* KO mice died within 8 days of birth (Fig. 2C, S2K), in line with a previous report^24^. Importantly, m^6^A, m^6,6^A, and i^6^A were highly elevated in the liver of *Adk* KO mice, while the adenosine level was comparable between WT and *Adk* KO mice (Fig. 2D). Taken together, our results demonstrate that one of the major roles of ADK is to metabolize m^6^A, m^6,6^A, and i^6^A once they are generated from RNA catabolism, and dysregulation of this pathway may lead to lethal consequences.

### Molecular insights into ADK-mediated m^6^AMP, m^6,6^AMP, and i^6^AMP formation

To confirm that m^6^A, m^6,6^A, and i^6^A are direct substrates of ADK, we performed *in vitro* kinase assays. ADK efficiently phosphorylated m^6^A, m^6,6^A, and i^6^A to their monophosphate forms *N*^6^-methlyadenosine monophosphate (m^6^AMP), *N*^6^,*N*^6^-dimethlyadenosine monophosphate (m^6,6^AMP), and *N*^6^-isopentenyladenosine monophosphate (i^6^AMP), respectively (Fig. 2E-F). The *k*_cat_/*K*_m_ values, which represents the catalytic efficiency^27, 28^ of m^6^A and i^6^A, were comparable to those of adenosine (m^6^A: 19 ± 2.3 mM^-1^ s^-1^, i^6^A: 18 ± 0.47 mM^-1^ s^-1^), and adenosine (19 ± 0.83 mM^-1^ s^-1^). Notably, the *k*_cat_/*K*_m_ value of m^6,6^A was even higher (44 ± 1.9 mM^-1^ s^-1^) than for other adenosines. ADK is reported to exhibit substrate inhibition by increasing the concentration of adenosine^29^. Indeed, in our experiments, adenosine exerted an inhibitory effect on ADK at concentrations higher than 20 µM. However, m^6^A, m^6,6^A, and i^6^A did not exhibit a substrate inhibition effect at any concentration tested (Fig. 2F). Thus, m^6^A, m^6,6^A, and i^6^A are readily phosphorylated to m^6^AMP, m^6,6^AMP, and i^6^AMP by ADK, respectively, as *bona fide* substrates. Consistent with these *in vitro* experiments, we found that exogenous addition of m^6^A, m^6,6^A, and i^6^A to cells led to the formation of m^6^AMP, m^6,6^AMP, and i^6^AMP, which were undetected in cells pre-treated with the ADK inhibitor (Fig. S2L).

Next, we performed *in silico* docking simulation based on the X-ray structure of human ADK complexed with adenosine (PDB: 1BX4)^30^ (Fig. 3A, S3A). The simulation revealed that similar to adenosine, m^6^A and m^6,6^A well fit in the substrate-binding site of ADK (Fig. 3B, S3B-C). Notably, the evolutionarily conserved L16, L40, L138, F201, A136, and F170 residues form a hydrophobic pocket, which seems to accommodate the *N*^6^ substituents of modified adenosines (Fig. 3C-F). i^6^A can also bind to the same binding site, but in a less optimal mode, suggesting that its binding may accompany the relocation or reorientation of the surrounding hydrophobic residues to accommodate the bulkier *N*^6^ substituent of i^6^A (Fig. S3D).

**Figure 3.**
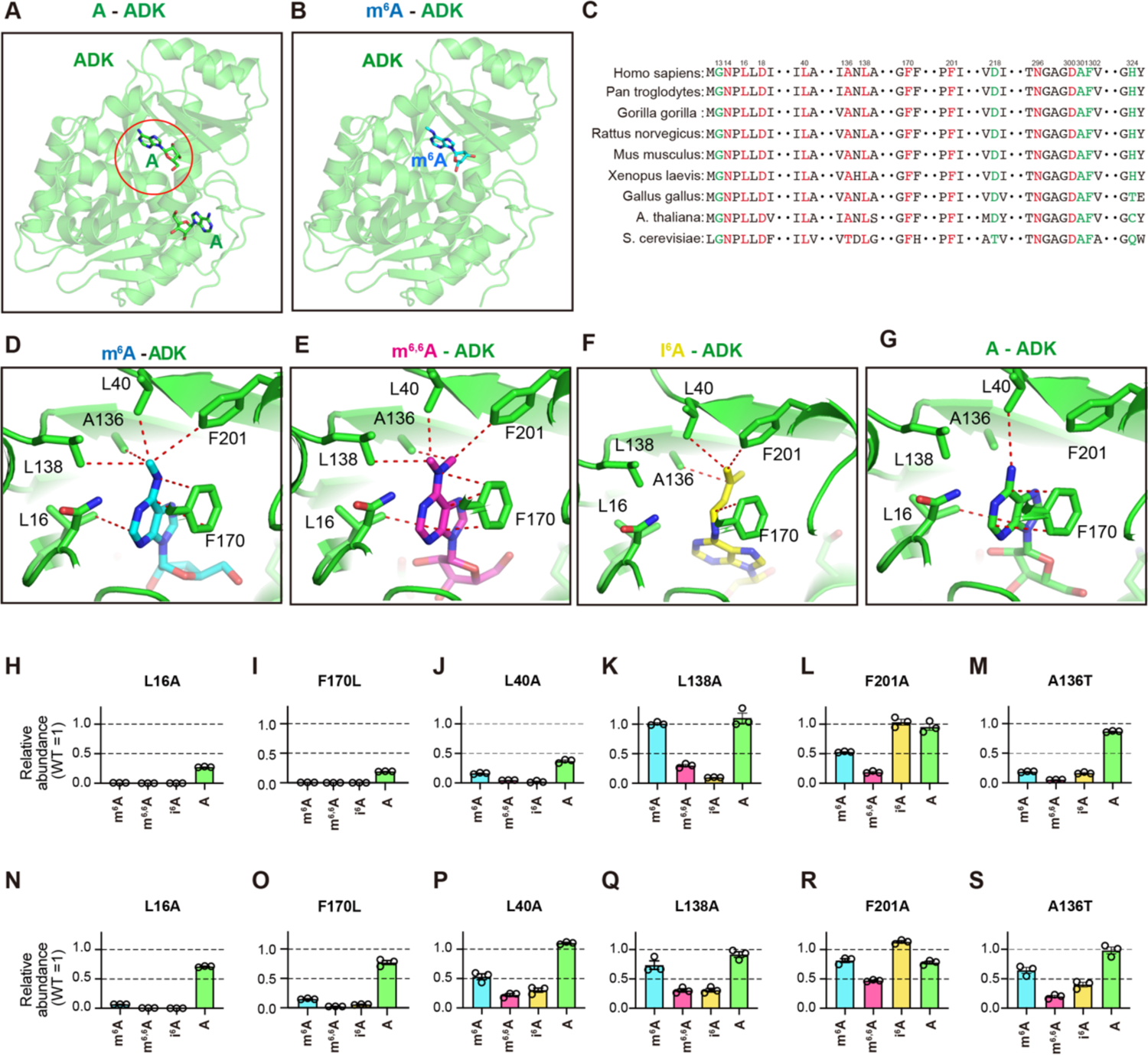
Docking of modified adenosines bound to ADK. (A) Crystal structure of ADK in complex with adenosine (PDB: 1BX4). Bound adenosine molecules are shown in a stick model. Carbon, oxygen and nitrogen atoms are colored in green, red and blue, respectively. The catalytic adenosine binding site of ADK is indicated by a red circle. (B) A docking model of m^6^A bound to ADK (cyan stick) (C) Cross-species alignment of ADKs: human (UniProt ID: P55263-2), chimpanzee (UniProt ID: A0A213TBB5), gorilla (UniProt ID: G3QRM8), rat (UniProt ID: Q64640), mouse (UniProt ID: P55264), frog (UniProt ID: Q6DJM1), chicken (UniProt ID: A0A8VOZQ37), *Arabidopsis thaliana* (UniProt ID: Q9SF85), and *Saccharomyces cerevisiae* (UniProt ID: P47143). Residues mutated in this experiment are colored red, and residues reported for human mutations are colored green. (D-G) Close-up views of the hydrophobic pocket of the (D) m^6^A-(cyan), (E) m^6,6^A-(magenta), (F) i^6^A-(yellow), or (G) adenosine (green)-bound ADK. Orange dashed lines represent van der Waals contacts. (H−S) Relative abundance of products (i.e., m^6^AMP for m^6^A, m^6,6^AMP for m^6,6^A, i^6^AMP for i^6^A, and AMP for adenosine) from *in vitro* assays (upper panel) or in cellular assays (lower panel) with each ADK mutant (compared to the amount of each product generated by WT ADK). The x-axis shows the substrates used.

To validate the predicted models, we generated recombinant ADK proteins with mutations at these residues and evaluated their kinase activities. The purine moiety of each adenosine molecule is sandwiched by L16 and F170 (Fig. 3D-G). Consistently, the mutation of L16A or F170L impaired the kinase activity toward both modified and unmodified adenosines (Fig. 3H-I). The L40A mutation also resulted in substantial loss of the kinase activity toward all modified adenosines examined, suggesting that L40A mutation distorts the hydrophobic pocket, hindering their binding to ADK (Fig. 3J). Some mutants exhibited selectivity for the modified adenosines: L138A mutant showed loss of activity to m^6,6^A and i^6^A but not to m^6^A and adenosine (Fig. 3K). This could be because L138A mutation may relocate L40 and A136, preventing their interactions with the dimethyl group of m^6,6^A or isopentenyl group of i^6^A (Fig. 3E-F). On the other hand, F201A mutation is likely to influence the relative positions of L138, L136 and L40, which may explain the loss of activity toward m^6^A and m^6,6^A, but not toward i^6^A and adenosine (Fig. 3L). A136 is highly conserved in eukaryotes except yeast with more hydrophilic threonine instead (Fig. 3C). Human ADK with A136T mutation showed large reduction in the kinase activity toward modified adenosines but not toward unmodified adenosine (Fig. 3M), highlighting the importance of the hydrophobic environment of the substrate-binding site in functional interaction with modified adenosines.

In addition to the mutations nearby the *N*^6^ modification site, we evaluated the mutations of residues interacting with the ribose moiety. The ribose moiety is predicted to locate at similar positions regardless of the *N*^6^ modifications (Fig.,S3A-D). D18A and D300A commonly abolished the kinase activity (Fig. S3E-F), because hydrogen bonds between D18 or D300 and the ribose moiety would be lost. The N296A mutant showed a substantial reduction of activity toward m^6,6^A and i^6^A (Fig. S3G), which may be explained by the closer location of the ribose O5’-hydroxyl group of these two modified adenosines to N296.

To further assess the effects of the mutations, we performed in cellular assay (Fig. 3N-S, S3H-J). On the whole, the intracellular kinase activities of the ADK mutants toward modified adenosines were compatible with the results of the *in vitro* assays. However, unlike the *in vitro* assays, ADK-mutant cells showed little difference in the level of unmodified AMP, possibly due to the *de novo* biosynthesis and salvage pathway for unmodified adenosine and AMP in cells.

Altogether, our results demonstrate that RNA catabolism-derived m^6^A, m^6,6^A and i^6^A are selectively recognized by ADK through hydrophobic interactions with specific residues, leading to their phosphorylation-dependent metabolism. These observations explain the low steady-state levels of intracellular m^6^A, m^6,6^A and i^6^A under normal conditions.

### ADAL-mediated deamination of m^6^AMP, m^6,6^AMP, and i^6^AMP to inosine monophosphate

Despite the high ADK activity for phosphorylating m^6^A, m^6,6^A, and i^6^A, steady-state levels of m^6^AMP, m^6,6^AMP, and i^6^AMP were very low or even undetectable in cells (Fig. 4A). This discrepancy indicates that m^6^AMP, m^6,6^AMP, and i^6^AMP are further subject to downstream metabolism. Indeed, when m^6^A, m^6,6^A, and i^6^A were added to cells, intracellular modified AMPs were only transiently formed and quickly disappeared from the cytosol (Fig. 4A). We found that metabolism of m^6^AMP, m^6,6^AMP, and i^6^AMP is not mediated by AMP deaminase, which is responsible for conversion of unmodified AMP to IMP (Fig. S4A).

**Figure 4.**
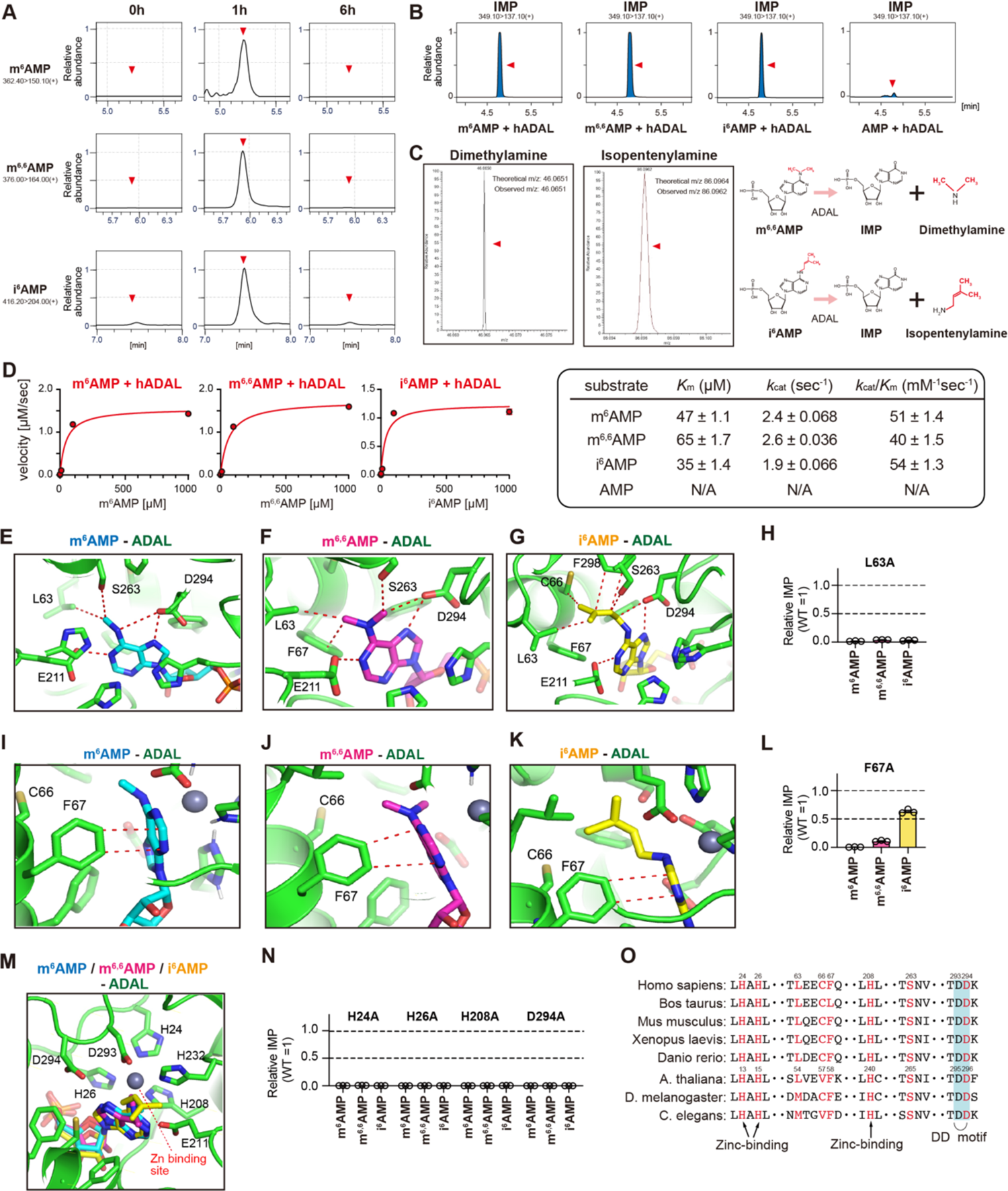
m^6^AMP, m^6,6^AMP, and i^6^AMP, but not AMP, are substrates for adenosine deaminase-like (ADAL). (A) Relative abundance of intracellular m^6^AMP, m^6,6^AMP, and i^6^AMP before, 1 h after, and 6 h after exogenous addition of m^6^A, m^6,6^A, and i^6^A in HEK293A cells. (B) Relative abundance of inosine monophosphate (IMP) measured by LC-MS following *in vitro* deamination assay using human ADAL protein. (C) Representative high-resolution mass spectra of dimethylamine (left) and isopentenylamine (right) resulting from removal of the dimethyl moiety from m^6,6^AMP or the isopentenyl moiety from i^6^AMP by ADAL, respectively, detected using an Orbitrap in positive ion mode. (D) Michaelis-Menten kinetic curves (left) and kinetic constants (right) of human ADAL for m^6^AMP, m^6,6^AMP, and i^6^AMP (n = 3). Results are means ± SEM. (E-G) Close-up views around the L63 and *N*^6^ modifications of (E) m^6^AMP (cyan), (F) m^6,6^AMP (magenta) and (G) i^6^AMP (yellow) in the docking model of modified AMPs bound to a homology model of human ADAL. Orange dashed lines represent van der Waals contacts or putative hydrogen bonds (D294, E211). (H) Relative abundance of IMP produced in *in vitro* deamination assays with ADAL L63A mutant (compared to the amount of IMP generated by WT ADAL). The x-axis shows the substrates used. (I-K) Close-up views around the F67 and *N*^6^ modifications of (I) m^6^AMP (cyan), (J) m^6,6^AMP (magenta) and (K) i^6^AMP (yellow). Orange dashed lines indicate Y-shaped χ-χ stacking interactions. (L) Relative abundance of IMP produced in *in vitro* deamination assays with the ADAL F67A mutant (compared to the amount of IMP generated by original ADAL). The x-axis shows the substrates used. (M) Close-up view around the zinc binding site including the ‘DD motif’ in modified AMPs-bound ADAL. (N) Relative abundance of IMP produced in *in vitro* deamination assays with ADAL H24A, H26A, H208A, and D294A mutants (compared to the amount of IMP generated by WT ADAL). The x-axis shows the substrates used. (O) Cross-species alignment of ADALs: human (UniProt ID: Q6DHV7), bovine (UniProt ID: Q0VC13), mouse (UniProt ID: Q80SY6), frog (UniProt ID: Q4V931), zebrafish (UniProt ID: Q4V9P6), *Arabidopsis thaliana* (UniProt ID: Q8LPL7), fruit fly (UniProt ID: Q9VHH7); *Caenorhabditis elegans* (UniProt ID: Q8IG39). Residues mutated in this experiment are colored red.

Recent studies reported that *Arabidopsis thaliana* and mammalian adenosine deaminase-like (*At*ADAL) can convert m^6^AMP to IMP^31,32^. However, whether this enzyme is also involved in the metabolism of other modified AMPs (m^6,6^AMP and i^6^AMP) is unknown. We performed *in vitro* deamination using recombinant *At*ADAL and human ADAL (*h*ADAL) with m^6^AMP, m^6,6^AMP, and i^6^AMP (Fig. S4B, 4B), and found that both *At*ADAL and *h*ADAL effectively converted m^6^AMP, m^6,6^AMP, and i^6^AMP to IMP. Notably, we successfully detected dimethylamine and isopentenylamine, reaction products alongside IMP upon deamination of m^6,6^AMP or i^6^AMP (Fig. 4C, S4C). We were not able to detect methylamine, the product of m^6^AMP deamination, due to instrumental limitations.

Detailed kinetics analysis revealed that all three modified AMPs showed comparable catalytic efficiency for ADAL (*k*_cat_/*K*_m_ values; m^6^AMP 51 ± 1.4 mM^-1^ s^-1^; m^6,6^AMP 40 ± 1.5 mM^-1^ s^-1^; i^6^AMP 54 ± 1.3 mM^-1^ s^-1^), while no valid values could be obtained with unmodified AMP (Fig. 4D). A previous report^32^ showed similar kinetic parameters of m^6^AMP to the present study. These results provide clear biochemical evidence that ADAL is responsible for the rapid clearance of m^6^AMP, m^6,6^AMP, and i^6^AMP by deamination to IMP.

### Molecular insights into ADAL-mediated deamination of modified AMPs

We next performed docking simulation and mutagenesis experiments to elucidate the molecular mechanism of ADAL-mediated deamination of modified AMPs. The structure prediction of *h*ADAL based on the crystal structure of m^6^AMP-bound *At*ADAL (PDB: 6ijn)^33^ showed the high similarity of these two structures (Fig. S4D), especially for the substrate-binding site (Fig. S4E). A catalytic zinc binding site is formed by the side chains of H24, H26, H208, and D293, which are highly conserved in the ADAL family^33,34^. The docking simulation revealed that modified AMPs (m^6^AMP, m^6,6^AMP, and i^6^AMP) fit snugly into the substrate-binding cavity of *h*ADAL, just as does m^6^AMP in *At*ADAL (Fig. S4F-H).

We found several key structural features that account for the preferred binding of modified AMPs to *h*ADAL. First, L63 and S263 locate close to the *N*^6^-substituent of the modified AMPs, forming van der Waals interactions (Fig. 4E−G). Second, F67 forms Y-shaped π-π stacking interactions with the adenine ring of the modified AMPs with distances of 3.5−4.5 Å (Fig. 4I−K). E211 and D294 seem to form hydrogen bonds with N atoms in the adenine ring of the modified AMPs (Fig. 4E-G). Additionally, F298 form van der Waals contact with the isopentenyl group of i^6^AMP (Fig. 4G)^33^. We also found that D294 next to D293 can be protonated due to the close proximity of acidic amino acids (Fig. 4M), likely leading to the increased pKa of D294. In this case, D294 seems likely to be hydrogen bonded to the *N*^7^ of the five-membered ring of the modified AMPs. We named this evolutionarily conserved motif as a ‘DD motif’ (Fig. 4M, 4O).

Consistent with the *in silico* modeling, the L63A mutation greatly abolished the potency of ADAL toward modified AMPs (Fig. 4H). The F67A mutation compromised the ADAL activity toward m^6^AMP and m^6,6^AMP, but to a lesser extent toward i^6^AMP (Fig. 4L). This could be because the bulkier modification of i^6^AMP allows its tight interactions with the surrounding residues even when the pocket is enlarged by the mutation (Fig. 4K). We also analyzed the effects of mutations of zinc-binding residues in ADAL (H24, H26, H208, D294, Fig. 4M). All ADAL mutants failed to deaminate modified AMPs (Fig. 4N). We also found that C66 might modulate the ADAL activity (Fig. S4J). In this context, C66 is substituted to valine in *C. elegans* and *A. thaliana* (Fig. 4O). Mutations of C66 to valine or other amino acids with bulkier side chains partially prevented the deamination of modified AMPs, suggesting that the bulky side chains might undergo steric hindrance with the *N*^6^ substituent to reduce the binding affinity of ADAL for modified AMPs.

Importantly, not only WT ADAL but also all ADAL mutant examined in this study failed to deaminate AMP to IMP (Fig. S4I, S4K-L), suggesting that ADAL is evolutionarily conserved to selectively metabolize RNA-derived m^6^AMP, m^6,6^AMP, and i^6^AMP (Fig. S4M).

### *In vivo* evidence for ADAL-mediated deamination of m^6^AMP, m^6,6^AMP, and i^6^AMP in *A. thaliana* and mammals

To explore the biological functions of ADAL, we generated ADAL KO HEK293 cells (Fig. S5A-C). As expected, m^6^AMP, m^6,6^AMP, and i^6^AMP, but not AMP, accumulated in the KO cells (Fig. 5A). We also observed an increase in extracellular m^6^A, m^6,6^A, and i^6^A, but not adenosine in the ADAL KO cells (Fig. S5D). In the human genome, four isoforms of ADAL have been reported (Fig. S5E). Among them, only the longest (full-length) isoform possesses zinc-binding H208 and DD motifs (residues 293−294), the putative catalytic sites essential for ADAL activity (Fig. 4N, 4O). These catalytic sites are well conserved in *AtA*DAL, although primary sequence conservation between *AtA*DAL and *h*ADAL is moderate (40% identity; Fig. S5E). As expected, only full-length *h*ADAL and *At*ADAL effectively suppressed the accumulation of the three modified AMPs when introduced into ADAL KO cells (Fig. 5B, S5F).

**Figure 5.**
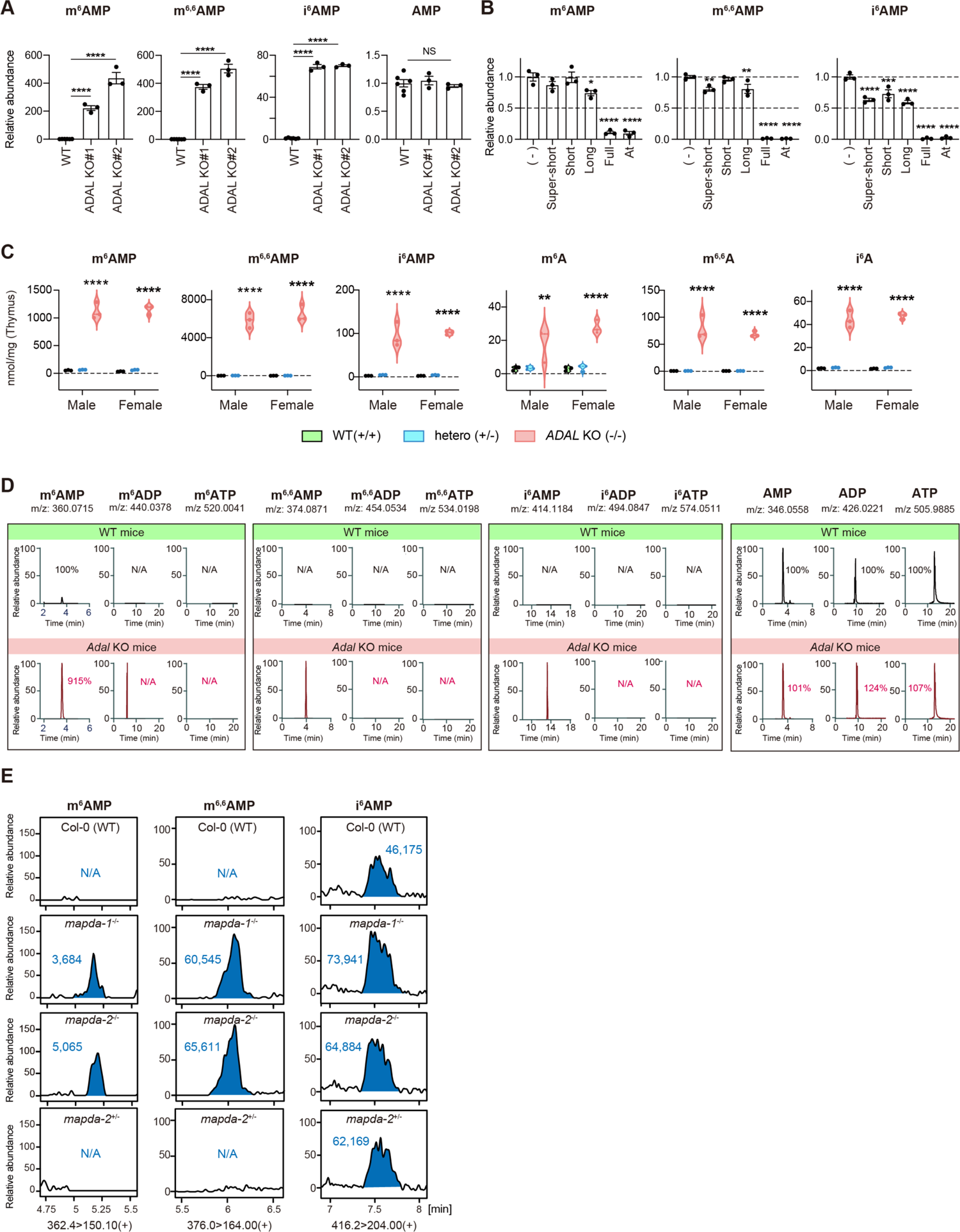
*In vivo* evidence of ADAL-mediated clearance of m^6^AMP, m^6,6^AMP, and i^6^AMP in mammals and *Arabidopsis thaliana*. (A) Relative abundance of intracellular m^6^AMP, m^6,6^AMP, i^6^AMP, and AMP in WT vs. two clones of ADAL KO cells (\*\*\*\**p* <0.0001, one-way ANOVA followed by Dunnett’s test) compared with the intracellular amount in WT cells. (B) Relative abundance of intracellular m^6^AMP, m^6,6^AMP, and i^6^AMP in ADAL KO cells (clone#1) and ADAL KO cells following reintroduction of each exogenous human ADAL isoform or *At-*ADAL. \**p* <0.05, \*\**p* <0.01, \*\*\**p* <0.001, and \*\*\*\**p* <0.0001 by one-way ANOVA followed by Dunnett’s test compared with ADAL KO cells without reintroduction. (C) Endogenous levels of modified AMPs (m^6^AMP, m^6,6^AMP, and i^6^AMP) and modified adenosines (m^6^A, m^6,6^A, and i^6^A) in the thymus of WT (black), heterozygous (blue), and *Adal* KO (red) mice. \*\**p* <0.01 and \*\*\*\**p* <0.0001 by two-way ANOVA followed by Šídák’s multiple comparison test. (D) Endogenous levels of modified AMPs, ADPs, and ATPs in the liver of *Adal* KO mice. While modified AMPs and unmodified AMP, ADP, and ATP were present, other modified ADPs or modified ATPs were not detected (peaks are all nonspecific). (E) Relative abundance of modified AMPs obtained from rosette leaf extracts of 1-month-old WT (Col-0), *mapda-1^-/-^*, *mapda-2^-/-^*, and *mapda-2^+/-^* Arabidopsis plants (MAPDA is the Arabidopsis ADAL orthologue).

To further analyze the functions of ADAL, we generated *Adal* KO mice (Fig. S5G, S5H), which were viable, fertile, and apparently normal (Fig. S5I, S5J). However, endogenous levels of both modified AMPs and modified adenosines were elevated in all organs examined in *Adal* KO mice (Fig. 5C, Table S2). The concentrations of the three modified AMPs varied between tissues, with thymus having the highest concentrations and muscle the lowest. Notably, among the three modified AMPs, m^6,6^AMP showed the highest concentration in all tissues (Table S2). Taken together, these results demonstrate that ADAL is responsible for the metabolism of m^6^AMP, m^6,6^AMP, and i^6^AMP *in vivo*.

Accumulation of m^6^AMP, m^6,6^AMP, and i^6^AMP in ADAL-deficient cells might lead to the formation of *N*^6^-modified ADP and ATP, which may interfere with RNA transcription. Adenylate kinase (AK) catalyzes the interconversion of various adenosine phosphates, including generating ADP from AMP and ATP as substrates^35^. When we performed *in vitro* assays by incubating unmodified AMP with AK and ATP, AMP was effectively converted to ADP (Fig. S5N). On the contrary, m^6^ADP/m^6^ATP, m^6,6^ADP/m^6,6^ATP, and i^6^ADP/i^6^ATP were barely detected when modified AMPs were incubated with AK and ATP *in vitro* (Fig. S5K−M). Most importantly, m^6^ADP/m^6^ATP, m^6,6^ADP/m^6,6^ATP, and i^6^ADP/i^6^ATP were below detectable levels in the liver of *ADAL* KO mice (Fig. 5D), suggesting that modified AMPs cannot be converted to diphosphate or triphosphate forms.

In addition to mammalian proteins, we isolated mutants of the *At*ADAL orthologue *N*^6^-methyl-AMP deaminase (*MAPDA*, At4g04480), including the previously-described *mapda-1* and *mapda-2*^31^, and a new *mapda-3* allele of the Nossen ectotype (Pst00481; Fig. S5O). Both m^6^AMP and m^6,6^AMP were undetectable in WT leaves but clearly detected in homozygous *mapda* mutants compared with cognate WT or heterozygous *mapda* plants (Fig. 5E, S5P). Furthermore, i^6^AMP, which was stably detected in WT leaves, was also elevated in *At*-ADAL-deficient leaves. The existence of i^6^AMP in WT leaves may reflect the fact that isoprenoid-type cytokinins including isopentenyladenine (i^6^Ade) are important cytokinin sources in plants, which can be biosynthesized by multiple enzymes^36^.

### Accumulation of modified AMPs dysregulate glucose metabolism in *Adal*

#### KO mice via AMP-activated protein kinase (AMPK) inhibition

To understand the pathophysiological role of ADAL-mediated deamination of m^6^AMP, m^6,6^AMP, and i^6^AMP, we performed a comprehensive phenotypic analysis in *Adal* KO mice. *Adal* KO mice exhibited similar characteristics to those of age-matched WT littermates, with no differences in histology (Fig. S6A), body weight (Fig. S6B), or general blood biochemistry related to liver and kidney functions (Fig. S6C-D, Table S3). Interestingly, *Adal* KO mice displayed significantly higher serum glycoalbumin (GA) and cholinesterase (ChE) levels than WT (Fig. 6A-B). To verify the impact of ADAL deficiency on glucose metabolism, we performed an intraperitoneal glucose tolerance test. In line with serum GA levels, *Adal* KO mice showed a mild but significant glucose intolerance compared with WT (Fig. 6C). We subsequently performed transcriptome analysis using liver and skeletal muscles, the tissues related to glucose metabolism. Gene ontology (GO) analysis revealed a significant (adjusted *p* <0.05) >2-fold change, indicating that the carbohydrate derivative metabolic process was commonly downregulated in the tested tissues (Fig. 6D-E, S6E). These molecular and physiological phenotypes suggest that accumulation of m^6^AMP, m^6,6^AMP, and i^6^AMP can lead to abnormal glucose metabolism.

**Figure 6.**
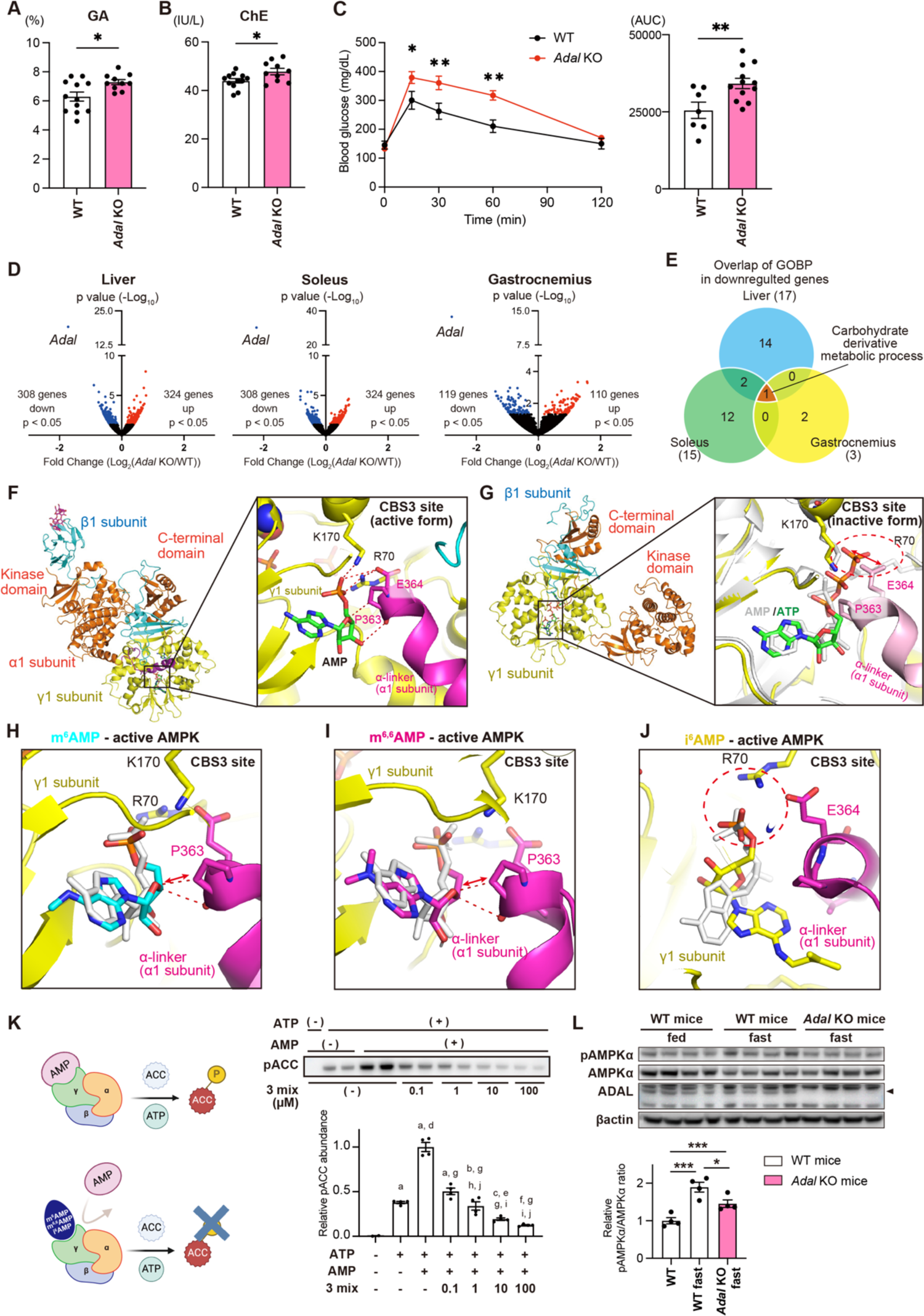
Modified AMPs allosterically inhibit AMPK activation and *Adal* KO mice exhibit dysregulated glucose metabolism. (A, B) Blood test results for serum glycoalbumin (A) and cholinesterase (B). n = 12 and n = 10 for WT and *Adal* KO mice, respectively. Results are means ± SEM. \**p* <0.05 between WT and *Adal* KO mice, by unpaired t-test. (C) (Left) Blood glucose levels during intraperitoneal glucose tolerance tests (IPGTT). n = 7 and n = 12 for WT and *Adal* KO mice, respectively. \**p* <0.05, \*\**p* < 0.01, two-way ANOVA followed by Šídák’s multiple comparison test. (Right) Area under the curve (AUC) during IPGTT. \*\**p* <0.01 by unpaired t-test. (D, E) Transcriptome analysis of tissues including liver, soleus muscle, and gastrocnemius muscle performed by RNA-seq in *ADAL* KO and WT mice at 30 weeks of age. (D) Volcano plot comparing gene expression in *ADAL* KO and WT mice. Blue and red dots indicate significantly altered genes in *ADAL* KO mice (>2-fold change, adjusted *p* <0.05 by DESeq2). (E) Venn diagram showing the overlap in gene ontology biological processes (GOBP) for downregulated genes between liver, soleus muscle, and gastrocnemius muscle. (F) Structure of the active form of the heterotrimeric human AMPK complex. (Left) The crystal structure of the phosphorylated human α1 β1 γ1 holo-AMPK complex bound to AMP (PDB, 4RER). The C-terminal and kinase domains of the α1 subunit are shown in orange, the α-linker in the α1 subunit in magenta, the β1 subunit in cyan, and the γ1 subunit in yellow. Three AMP molecules are bound to Cystathionine β-synthetase (CBS) sites. (Right) Close-up view of the CBS3 site in the activated form, in which the α phosphate group of AMP (shown in orange) is recognized by the R70 and K170 residues of the γ1 subunit (red dashed lines), and the α-linker interacts with the γ1 subunit through the electrostatic interaction between the E364 residue of the α-linker and the R70 residue of the γ1 subunit (red dashed lines). (G) Structure of the inactive form of the heterotrimeric human AMPK complex. (Left) The cryo-EM structure of ATP-bound, inactive AMPK (PDB, 4JHj, Fab and maltose-binding protein are omitted for clarity). ATP is bound to CBS3. (Right) Close-up view of the CBS3 site in the inactivated form. The active-form AMPK structure is superimposed onto the γ1 subunit, The γ1 subunit and AMP of the active form are shown in light gray, and the α-linker in light pink. A red dashed circle shows the steric hindrance caused by the bulky β- and γ-phosphate groups of ATP (shown by the red double-headed arrows), which disrupts the interaction between E364/P363 of the α-linker and K170/R70 of the γ-subunit and causes the kinase domain to be largely displaced from the orthosteric active state.. (H-J) Docking simulation of (H) m^6^AMP (cyan), (I) m^6,6^AMP (magenta), and (J) i^6^AMP (yellow) bound to active-form AMPK. Unmodified AMP is shown in light gray. Red two-headed arrows in (H) and (I) indicate the repulsion between the P363 residue of the α-linker and the ribose moiety of m^6^AMP or m^6,6^AMP. The red dashed circle in (J) highlights the conformational change from the orthosteric active state. (K) *In vitro* phosphorylation assay to determine AMPK activity. (Left) Schematic diagram of the assay. A truncated acetyl-CoA carboxylase 1 (ACC), human recombinant AMPK, AMP (1 µM), and with (bottom) or without (top) modified AMPs were incubated *in vitro* in the presence of ATP, and the phosphorylated ACC was detected. (Right, upper panel) SDS-PAGE of phosphorylated ACC. (Right, lower panel) Densitometry results. a, *p* <0.0001 vs. control; b, *p* <0.001 vs. control; c, *p* <0.05 vs. control; d, *p* <0.0001 vs. ATP(+); e, *p* <0.01 vs. ATP(+); f, *p* <0.001 vs. ATP(+); g, *p* <0.0001 vs. ATP(+)AMP(+); h, *p* <0.05 vs. ATP(+)AMP(+)3mix(0.1); i, *p* <0.0001 vs. ATP(+)AMP(+)3mix(0.1); j, *p* <0.01 vs. ATP(+)AMP(+)3mix(1), by one-way ANOVA followed by Tukey’s multiple comparison test. (L) Representative western blotting results for phosphorylation of AMPKα after 16 h of fasting in WT vs. *Adal* KO mice (n = 4 each, upper panel). Densitometry results are shown in the lower panel. \**p* <0.05 and \*\*\**p* <0.001 by one-way ANOVA followed by Tukey’s multiple comparison test. Results are means ± SEM.

The altered glucose metabolism prompted us to examine the effect of modified AMPs on AMPK, given the essential role of AMPK in energy sensing and metabolism in tissues^37^. AMP activates AMPK by competitively binding to cystathionine beta-synthase (CBS) 3 site within the γ subunit of AMPK^38, 39, 40, 41, 42, 43, 44^. In the active form of AMPK, the α-linker in the α1 subunit associates with γ subunit, in which AMP contributes to their interface (Fig. 6F). In contrast, when ATP is bound to CBS3, the bulky β- and γ-phosphate groups of ATP would undergo steric hindrances with E364 and P363 in the α-linker, leading to large displacement of the α-linker and kinase domain, followed by inactivation of AMPK (Fig. 6G).

Given the structural resemblance between modified AMPs and AMP, we surmise that elevated m^6^AMP, m^6,6^AMP, and i^6^AMP might be implicated in the regulation of AMPK activity. Indeed, docking simulation of m^6^AMP, m^6,6^AMP, and i^6^AMP revealed that the modified AMPs would bind to the CBS3 site in the γ subunit of AMPK over AMP (Fig. 6H-J). The ribose moiety of m^6^AMP or m^6,6^AMP is repelled by P363 due to steric repulsion, suggesting that the interaction between the α linker and the γ1 subunit might be weakened, thereby possibly suppressing the AMPK activation (Fig. 6H-I). In the i^6^AMP-bound model, although no steric repulsions are observed, the alteration in the coordination of the phosphate group or the elimination of the interaction between the ribose moiety and P363 may lead to the compromised interaction between the α linker and γ1 subunit (Fig. 6J). Docking simulations also showed that m^6^AMP and i^6^AMP may bind to the ATP-binding site in the kinase domain of the α-subunit in the active form of AMPK, thereby competing with its substrate ATP (Fig. S6F). These observations suggest that the modified AMPs allosterically inhibit AMP-mediated AMPK activation by competing with AMP for binding to the CBS3 site, and possibly prevent its kinase activity by competing with ATP for binding to the kinase domain.

To verify the validity of the above docking simulation, we performed *in vitro* AMPK kinase assays using m^6^AMP, m^6,6^AMP, and i^6^AMP as substrates. Compared with unmodified AMP, m^6^AMP, m^6,6^AMP, or i^6^AMP alone showed a very weak potency to activate AMPK (Fig. S6G). Instead, we observed that m^6,6^AMP and i^6^AMP effectively inhibited AMP-mediated AMPK activation, with the inhibitory effect of m^6^AMP being moderate (Fig. S6H). We then performed a kinase assay using the mixture of modified AMPs to recapitulate intracellular conditions. The modified AMPs dose-dependently suppressed AMPK activity *in vitro* (Fig. 6K). Importantly, the modified AMPs had an inhibitory effect even at 100 nM, which could be achieved in *Adal* KO mouse tissues (Fig. 5C).

Furthermore, we verified the inhibitory effect of modified AMPs in ADAL KO cells and *Adal* KO mice. In line with the *in vitro* study, starvation-induced AMPK activation was significantly suppressed in ADAL KO cells (Fig. S6I). Likewise, phosphorylation of AMPK was significantly suppressed in the liver of fasted *Adal* KO mice, indicating that AMPK activation was significantly suppressed (Fig. 6L).

Thus, once m^6^A, m^6,6^A, and i^6^A are phosphorylated to their monophosphate forms by ADK, m^6^AMP, m^6,6^AMP, and i^6^AMP are quickly metabolized to IMP by ADAL to prevent their potential influence on AMPK activation (Fig. S6J). It should be noted that although pathological mutations of ADAL have not been reported in humans, many missense variants at the catalytically critical amino acids mediating deamination of modified AMPs, such as H24, H26, L63, C66, H208, S263, and D294, have been reported in the large population genome database (Table S4). Thus, aberrant accumulation of m^6^AMP, m^6,6^AMP, and i^6^AMP may be linked to human diseases, especially metabolic diseases such as diabetes.

#### Disease-related ADK mutations and links to m^6^A, m^6,6^A, and i^6^A

Finally, we explored the pathophysiological importance of ADK-mediated metabolism of modified adenosines. Patients carrying pathological mutations, including G13E, D218A, A301E, F302S, and H324R in the *ADK* gene^45,46^, develop a clinical phenotype including hepatic and neurologic impairment^46^. The molecular pathogenesis has been attributed to methionine metabolism^45^, but direct evidence remains elusive. Thus, we generated recombinant ADK proteins with these pathogenic mutations and evaluated their kinase activities.

The *in vitro* kinase assays showed that G13E, D218A, A301E, and F302S are complete loss-of-function mutations while the H324R mutation is a partial loss-of-function mutation to both modified and unmodified AMPs (Fig. 7A). Next, we introduced the *ADK* mutants into ADK KO cells and examined the levels of modified and unmodified adenosines. In line with the *in vitro* assays, ADK KO cells expressing G13E, D218A, A301E, and F302S mutants accumulated more m^6^A, m^6,6^A, and i^6^A in the supernatant than WT ADK, but levels of unmodified adenosine remained unchanged (Fig. 7B). Accordingly, formation of modified AMPs was drastically attenuated in cells expressing G13E, D218A, A301E, and F302S mutants, while levels of unmodified AMP levels were unchanged compared with WT cells (Fig. 7C). On the other hand, cells expressing the H324R mutant showed moderate accumulation of modified adenosines extracellularly, and modified AMPs intracellularly, in accordance with the *in vitro* kinase assay results. Collectively, disease-related ADK mutants (G13E, D218A, A301E, F302S, and H324R) are strongly associated with aberrant modified adenosine metabolism.

**Figure 7.**
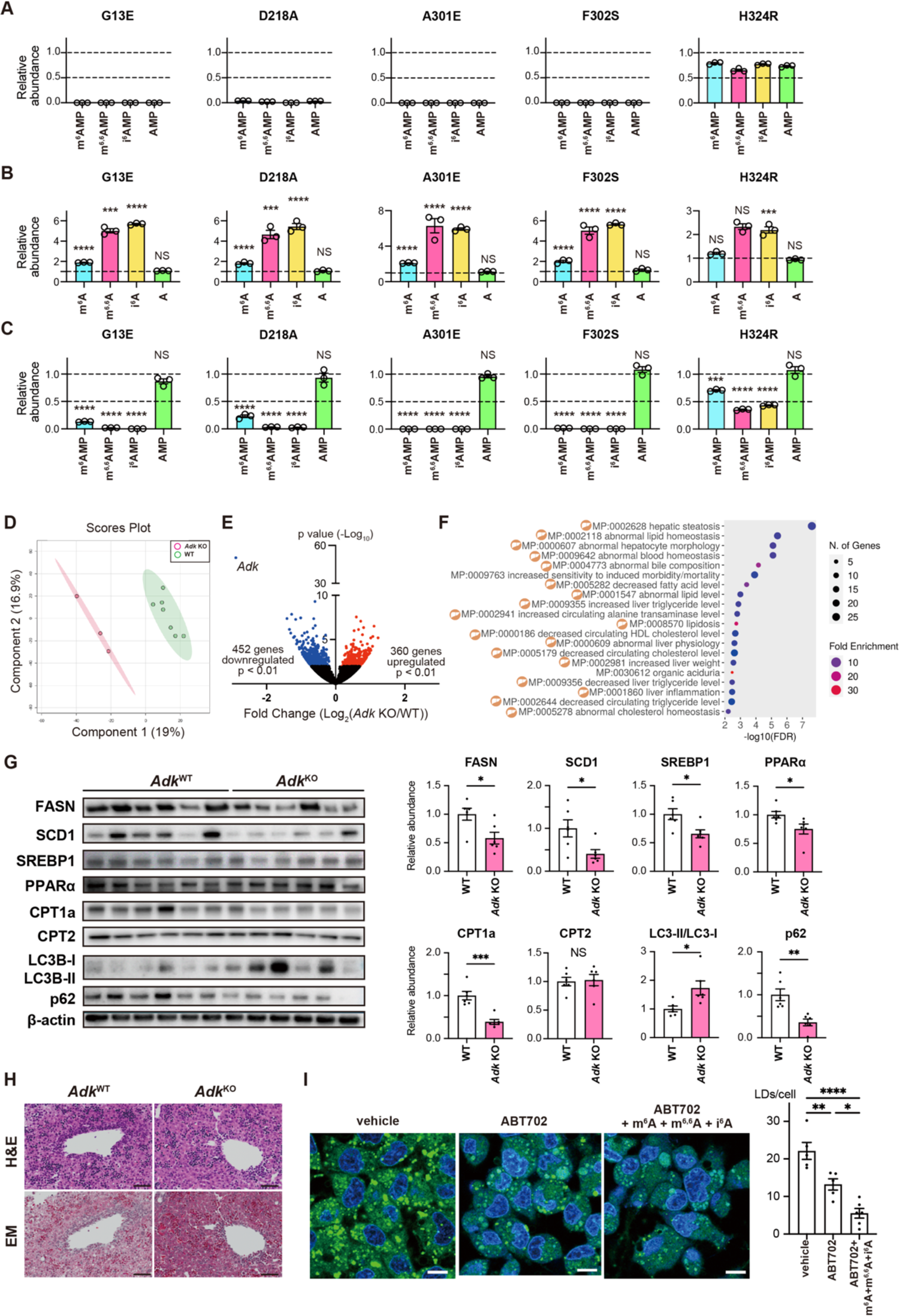
ADK deficiency causes a massive increase in modified adenosines, which leads to defective lipid metabolism. (A) Relative abundance of products (i.e., m^6^AMP for m^6^A, m^6,6^AMP for m^6,6^A, i^6^AMP for i^6^A, and AMP for adenosine) from *in vitro* assays with each human pathogenic ADK mutant (compared to the amount of each product generated by WT ADK). (B, C) Relative abundance of m^6^A, m^6,6^A, i^6^A, and adenosine in the supernatant (B) or corresponding m^6^AMP, m^6,6^AMP, i^6^AMP, and AMP inside cells (C) when ADK encoding each human pathogenic mutation is reintroduced into ADK KO cells. The abundance was compared to that of each nucleoside/nucleotide generated by WT ADK. \*\*\**p* <0.001 and \*\*\*\**p* <0.0001 by one-way ANOVA followed by Dunnett’s test, n = 3. (D) Principal component analysis of non-target metabolomics between the liver of WT (green, n = 7) and *Adk* KO mice (pink, n = 3) at postnatal day 5. (E) Volcano plot from RNA-seq analysis comparing gene expression in the liver of *Adk* KO and WT mice (>2-fold change, adjusted *p* <0.01 by DESeq2). (F) The top 20 enrichment pathways of downregulated genes using GO. Dot size represents the number of enriched genes and dot color represents the fold enrichment. The pathways involved in liver and/or lipid metabolism are marked with a liver symbol. (G) (Left) Western blotting analysis of the levels of proteins related to lipid metabolism or autophagy in the liver of WT (n = 6) and *Adk* KO (n = 6) mice. (Right) Densitometry results. \**p* <0.05, \*\**p* <0.01, and \*\*\**p* <0.001 between WT and *Adk* KO mice, by unpaired t-test. Results are means ± SEM. (H) Representative H&E staining and Elasitica Masson staining of liver in WT and *Adk* KO mice. Scale bar = 50 µm. (Left) Representative BODIPY staining of lipid droplets (LDs, green) and DAPI (blue) in HepG2 cells. HepG2 cells were incubated with ABT702 (10 µM) for 1 h, followed by separate addition of the three modified adenosine mixtures (100 nM of m^6^A, m^6,6^A, and i^6^A) for 24 h. Scale bar = 10 µm. (Right) LD counts were analyzed using ImageJ software. \**p* <0.05, \*\**p* <0.01, and \*\*\*\**p* <0.0001, by one-way ANOVA followed by Tukey’s multiple comparison test. Results are means ± SEM.

#### Molecular pathogenesis of ADK deficiency

Finally, we sought to understand the impact of the accumulation of modified adenosines *in vivo* and its relevance in ADK deficiency. Hepatic impairment is the predominant symptom in human ADK deficiency^46^. Moreover, *Adk* is ubiquitously expressed in mouse tissues with the highest expression in liver (Fig. S7A). We performed untargeted and targeted metabolomics using postnatal liver tissues of *Adk* KO mice before their death. Principal component analysis of untargeted metabolomics data revealed that the metabolic profile of the liver was largely different between WT and *Adk* KO mice (Fig. 7D). Subsequent targeted metabolomics analysis of the nucleotide pool revealed that nucleotides, including phosphoribosyl pyrophosphatase (PRPP) and cytidine triphosphatase (CTP), were upregulated in the liver of *Adk* KO mice (Fig. S7B-C). Importantly, methionine levels and methionine cycle-related metabolites were unchanged in the liver of KO mice (Fig. S7D), suggesting that the severe clinical phenotypes of ADK deficiency were not caused by the proposed abnormal methionine metabolism.

Subsequent transcriptome analysis of the livers of WT and *Adk* KO mice and GO analysis revealed that most of the downregulated genes in *Adk* KO mouse liver were significantly correlated with lipid metabolism (Fig. 7E-F), while upregulated genes were related to stress responses (Fig. S7E). We also performed transcriptome analysis in the brain of *Adk* KO mice, but gene expression changes in brain were more subtle than those in liver (Fig. S7F). Nevertheless, there were several genes related to lipid metabolism that were commonly decreased in both liver and brain (Fig. S7G-H).

We then validated the results of the transcriptome analysis by assessing the levels of proteins related to lipid metabolism. Lipid metabolism consists of a *de novo* synthetic pathway (lipogenesis) and a catabolic process leading to β-oxidation^47^. Consistent with the transcriptome analysis, proteins related to lipogenesis, including fatty acid synthase (FASN) and stearoyl-CoA desaturase-1 (SCD1), and catabolic processes including carnitine palmitoyltransferase 1 (CPT1), were significantly suppressed in the liver of *Adk* KO mice (Fig. 7G). Sterol regulatory element-binding protein1 (SREBP1) and peroxisome proliferator-activated receptorα (PPARα), key transcription factors integrating lipid metabolism^47,48^, were both significantly downregulated (Fig. 7G), indicating that the major lipid metabolism process is globally impaired in the liver of *Adk* KO mice. Importantly, histological analysis of *Adk* KO liver revealed that the lipid droplet (LD) content in *Adk* KO mice was lower than in WT mice (Fig. 7H). As lipids are the major energy source for cells, defective lipid metabolism can trigger stress responses including autophagy^49, 50^. Indeed, we observed an increase in autophagy-related signatures characterized by an increased LC3-II/LC3-I ratio (LC3 lipidation) and a concomitant decrease in p62 in the liver of *Adk* KO mice (Fig. 7G). Finally, we sought to elucidate whether elevated modified adenosines are sufficient for abnormal lipid metabolism. To this end, we treated HepG2 cells with modified adenosines and visualized cytoplasmic LDs, which are crucial organelles for energy storage and lipid homeostasis, and are degraded by autophagy^51^. As expected, LD formation was impaired upon ADK inhibition and, importantly, further inhibited upon exogenous addition of the mixture of modified adenosines (Fig. 7I). Lastly, consistent with these findings, proteins related to lipid metabolism were significantly downregulated and proteins related to autophagy responses were significantly upregulated in HepG2 cells treated with ADK inhibitor and modified adenosines (Fig. S7I-J).

Taken together, these results demonstrate that ADK serves as a first-line responder to RNA-derived m^6^A, m^6,6^A, and i^6^A through phosphorylation of these modified adenosines, followed by ADAL-mediated deamination to IMP. Additionally, they indicate that the molecular pathology of ADK deficiency is associated with the aberrant accumulation of cytotoxic modified adenosines, which impairs lipid metabolism and ultimately leads to the onset of catastrophic symptoms (Fig. S7K).

## DISCUSSION

### Biological implications of the ADK-ADAL pathway

Diverse RNA modifications have been implicated in post-transcriptional gene expression regulation and subsequent complex biological phenomena^1,2^. RNA catabolism yields numerous modified nucleosides both intracellularly and extracellularly. However, the biology of these modified nucleosides has remained elusive. In this study, we investigated the metabolism and pathophysiological impacts of RNA-derived modified nucleosides in cells and animals. In this study, we show that RNA-derived modified nucleosides have distinct biological properties and are subjected to differential metabolic regulation. While most RNA-derived modified nucleosides are inert and stable, several modified nucleosides including Am, m^7^G, m^6^A, m^6,6^A, and i^6^A are rapidly metabolized (Fig. 1D). Among these unstable nucleosides, Am and m^7^G are harmless to cells or even enhance cell viability in some cell lines (Fig. S1A-G). By contrast, m^6^A, m^6,6^A, and i^6^A exhibit toxicity toward a broad range of cell lines (Fig. 1A-B, Fig. S1A-H). Interestingly, metabolism of m^6^A, m^6,6^A, and i^6^A is tightly regulated in a stepwise manner. These modified adenosines are first phosphorylated by ADK to their monophosphate forms (m^6^AMP, m^6,6^AMP, and i^6^AMP), followed by ADAL-mediated deamination to IMP (Fig. S4M). Accordingly, ADK deficiency induced the accumulation of m^6^A, m^6,6^A, and i^6^A in cells and mice (Fig. 2A, 2D), and ADAL-deficient cells, mice, and plants accumulated m^6^AMP, m^6,6^AMP, and i^6^AMP (Fig. 5A, 5C, and 5E). The ADK-ADAL pathway thus explains why steady-state levels of m^6^A, m^6,6^A, and i^6^A, as well as their monophosphate forms, are normally very low or even undetectable in cells and tissues compared with other modified nucleosides. Furthermore, we show that ADK deficiency and ADAL deficiency induced lethal and adverse phenotypes, respectively, in mice. Thus, the ADK-ADAL pathway is a defense system against m^6^A, m^6,6^A, and i^6^A, the three cytotoxic products of RNA catabolism. In addition, we previously showed that RNA catabolism-derived extracellular m^6^A is a signaling molecule that activates the adenosine A3 receptor and induces allergic reaction and inflammation^21^. Thus, it is conceivable that ADK-mediated m^6^A metabolism also serves as a negative feedback mechanism to inactivate the A3 receptor and subsequent signaling transduction which triggers immune responses.

### ADK-mediated phosphorylation of modified adenosines

Unmodified adenosine is a substrate of ADK, and the resulting product AMP is the building block of nucleic acids, a crucial energy system component, and a signaling molecule. Our study provides biochemical, structural, and *in vivo* evidence that m^6^A, m^6,6^A, and i^6^A, which are modified adenosines derived from RNA catabolism, are also the endogenous substrates of ADK. m^6^A is one of the most abundant RNA modifications found on mRNA, rRNA, and non-coding RNA^52^. Extensive studies on m^6^A in recent years identified its intracellular role in RNA stability, translation, and microRNA biogenesis^14, 53, 54^, but the extracellular role of m^6^A has been elusive. The presence of m^6^AMP was reported in plants and mammals^31,55,56^, but it was simply considered an RNA degradation product. Since lysosome is one of the major RNA-degrading organelles, m^6^AMP will be quickly dephosphorylated to m^6^A by lysosomal phosphatase, and since m^6^AMP cannot be formed in ADK-deficient cells, it is likely that most intracellular m^6^AMP is generated by ADK-mediated phosphorylation of RNA catabolism-derived m^6^A. In addition to m^6^A, m^6,6^A is among the most abundant and evolutionarily conserved modifications, and is exclusively present in rRNA. Formation of m^6,6^A is catalyzed by DIMT1L in mammals, and it plays a key role in ribosome biogenesis^57,58^. However, neither the metabolic pathway of m^6,6^A nor its pathophysiological impacts as a modified nucleoside have been reported. In the present study, we show that rRNA catabolism-derived m^6,6^A possesses cytotoxicity and is rapidly phosphorylated by ADK. To the best of our knowledge, this is the first report identifying m^6^A and m^6,6^A as the direct substrates of ADK.

Compared with m^6^A and m^6,6^A, i^6^A and its derivatives have long been studied as cytokinins in the field of plant hormones, which are essential for plant cell growth and differentiation as activators of cytokinin receptors^36,59^. In mammals, tRNA isopentenyltransferase 1 (TRIT1) is responsible for i^6^A modification at position 37A of a subset of cytosolic and mitochondrial tRNAs^60^. i^6^A modification is essential for reading frame maintenance, and dysregulation of this modification has been implicated in the development of mitochondrial diseases^60^. Unlike in plants, the nucleoside form of i^6^A does not exert a growth hormone-like effect in mammals because cytokine receptors are not conversed. To date, all studies on i^6^A in mammals have been performed using synthetic i^6^A, specifically on human cancer cells and primary cells, and the metabolic regulation of endogenous i^6^A and its physiological implications remain elusive. In line with previous studies^61, 61, 62^, we show that exogenously added i^6^A had an adverse effect on cell culture (Fig. S1H), and it could be phosphorylated to i^6^AMP by ADK (Fig. S2L). Importantly, using ADK KO cells and mice, we showed that endogenous i^6^A can be substrates of ADK (Fig. 2B and 2D).

Taken together, our results demonstrate that once m^6^A, m^6,6^A, and i^6^A are generated by RNA catabolism, they are quickly recognized and phosphorylated by ADK as endogenous substrates. This phosphorylation is indispensable to prevent their accumulation and subsequent cytotoxicity.

### Pathogenic ADK mutations and modified adenosines

One of the most important findings in this study is the pathophysiological link between the modified adenosines and human disease. Missense mutations in the *ADK* gene, including G13E, D218A, A301E, F302S, and H324R, have been reported in patients with liver disease, dysmorphic features, epilepsy, and developmental delay^45,46^, and there is no curative treatment^45^. ADK deficiency patients harboring G13E, D218A, A301E, and F302S mutations are often diagnosed at an early age with severe symptoms, whereas patients with the H324R mutation are typically diagnosed at ≥20 years of age and their symptoms are mild^46^. In the present study, using *in vitro* kinase assays, we show that G13E, D218A, A301E, and F302S are complete loss-of-function mutations while H324R is a partial loss-of-function mutation associated with the three modified adenosines (Fig. 7A). Furthermore, cells expressing pathogenic ADK mutants accumulated extracellular modified adenosines and the concomitant loss of intracellular modified AMP (Fig. 7B-C). Combining clinical information and our results, the ADK deficiency phenotype appears to correlate well with the ability of ADK to phosphorylate modified adenosines. Mechanistically, residues F302 and H324 are predicted to interact with the ribose of ATP, while A301 is positioned next to D300 that interacts through hydrogen bonding with the ribose of adenosine^30^. Thus, mutations of these three residues in ADK are likely to affect kinase activity regardless of modifications at the *N*^6^ position. On the other hand, G13 and D218 are not engaged in direct binding with substrates, and the reason why these two mutations are loss-of-function remains to be elucidated. It should be noted that although the disease-related ADK mutants are also loss-of-function for unmodified adenosine *in vitro*, steady-state levels of intracellular adenosine and AMP remained unchanged in mutant cells compared with WT cells. This is because besides ADK, adenosine and AMP can be formed through multiple pathways, including *de novo* purine biosynthesis and salvage pathways. These results strongly suggest that accumulation of modified adenosines contributes significantly to the pathogenesis of ADK deficiency. Furthermore, given that modified adenosines can only be phosphorylated and metabolized by the ADK pathway, and their robust accumulation upon ADK deficiency, m^6^A, m^6,6^A, and i^6^A are the most suitable metabolites for the diagnosis of ADK deficiency.

### Molecular pathogenesis of ADK deficiency and modified adenosines

The pathology of ADK deficiency has been attributed to the aberrant elevation of methionine^45^. Because adenosine is one of the reaction products of the methionine cycle, it has been proposed that ADK deficiency can elevate intracellular adenosine levels, leading to slowdown of the methionine cycle and build-up of methionine. However, some ADK deficiency patients showed no hypermethioninemia or even spontaneous declining methionine levels over years without phenotype withdrawal^63^. Furthermore, the clinical effects of a methionine-restricted diet appear to be limited. Thus, the involvement of methionine in ADK deficiency is not as direct as in other hypermethioninemia diseases. Supporting this view, we found that methionine and other metabolites in the methionine and folate cycles did not differ between WT and *Adk* KO mice (Fig. S7D). Interestingly, our study suggests that aberrant lipid metabolism may be involved in the pathogenesis of ADK deficiency. Transcriptome and subsequent protein expression analysis by western blotting revealed that lipogenesis and downstream lipid metabolism are globally impaired in *Adk* KO mice (Fig. 7E−G).

Furthermore, the formation of LDs, whose primary function is energy storage and provision, is decreased under ADK inhibition (Fig. 7I), consistent with previous results from experiments involving human liver biopsy and mice with hepatocyte-specific ADK disruption or overexpression^64^. Most importantly, LD formation was further decreased by exogenous addition of a mixture of modified adenosines (m^6^A, m^6,6^A, and i^6^A; Fig. 7I), which can be considered chronic inhibition of ADK as occurs in *Adk* KO mice or ADK-deficient patients, suggesting that ADK deficiency exacerbates the effects of m^6^A, m^6,6^A, and i^6^A. In general, the liver is highly dependent on lipids as an energy source from the neonatal stage through childhood^65^, especially under stressful conditions such as starvation or infection. Thus, our results suggest that elevated modified adenosines might contribute to the pathogenesis of ADK deficiency through impairment of hepatic lipid metabolism.

### Molecular and physiological functions of ADAL

Our study illuminated the metabolic pathway of endogenous modified adenosines by elucidating the molecular functions of ADAL associated with m^6^AMP, m^6,6^AMP, and i^6^AMP generated by ADK. ADAL was first biochemically purified from rat liver homogenate as an enzyme possessing deamination activity for m^6^AMP^66^, and it was named *N*^6^-Methyl-AMP aminohydrolase^66^. Subsequent studies found that ADAL is widely conserved in eukaryotes^67^. To date, most studies have focused on the biochemical role on m^6^AMP. In the present study, we expand the role of ADAL on the endogenous modified AMPs and provide biochemical and *in vivo* evidence that ADAL is responsible for the deamination of m^6^AMP, m^6,6^AMP, and i^6^AMP (Fig. 5C). The deaminase activity of ADAL is highly specific to m^6^AMP, m^6,6^AMP, and i^6^AMP, but not to unmodified AMP (Fig. 4B). Our homology modeling provides an explanation for the substrate specificity. First, the methyl and isopentenyl groups of the three modified AMPs, which are not present in unmodified AMP, engage in hydrophobic interactions with lipophilic amino acids of ADAL including L63 and F67. In addition, D294 of ADAL forms hydrogen bonds with the amino group of unmodified AMP, which pulls AMP away from the optimal position for nucleophilic attack and subsequent deamination (Fig. S4I and S4L).

Why m^6^AMP, m^6,6^AMP, and i^6^AMP need to be deaminated by ADAL *in vivo* has not been investigated. Moderate impairment of root growth was observed in *At*ADAL-deficient *A. thaliana* plants^31^, but the mechanism is unclear. In the present study, we generated *Adal* KO mice and found that accumulation of modified AMPs is associated with aberrant glucose metabolism (Fig. 6A-E).

Furthermore, we found that the modified AMPs can allosterically inhibit AMPK at physiologically relevant concentrations^68^. AMPK is one of the master regulators of energy metabolism through competitive binding of AMP, ADP, and ATP to three sites in its γ subunit, of which cystathionine beta-synthase domain 3 (CBS3) is the primary sensor of AMP^40,41^. AMPK is dysregulated in several chronic diseases including diabetes, obesity, inflammation, and cancer^69^. Structural modeling suggests that the modified AMPs binds to the γ subunit, thereby inhibiting AMPK activation (Fig. 6H−J), and this was supported by the *in vitro* AMPK assay results (Fig. 6K). Importantly, using ADAL*-*deficient mice, we demonstrated the inhibitory effects of m^6^AMP, m^6,6^AMP, and i^6^AMP on physiological stress-induced AMPK activation (Fig. 6L), which might explain the glucose intolerance observed in *Adal* KO mice. Taken together, these results suggest that one of the physiological roles of ADAL, at least in mammals, is to metabolize the modified AMPs generated by ADK to prevent their antagonistic effects on AMP-regulated proteins, such as AMPK. Interestingly, Pisanti *et al*. reported that exogenously added i^6^A might induce AMPK activation through i^6^AMP in cell culture^62^. However, given the substantial concentration of exogenous i^6^A and the lack of *in vitro* kinase assay and *in vivo* evidence in their study, the activation effect of exogeneous i^6^A might be secondary under non-physiological conditions.

Recently, murine ADAL was shown to catalyze deamination of DNA-derived 2-deoxy-*N*^6^-methyladenosine monophosphate (6mdAMP), which might potentially induce aberrant epigenetic markers via modified *N*^6^-methyadenosine di- or triphosphoate^56^. Using *in vitro* kinase assays, we showed that m^6^AMP, m^6,6^AMP, and i^6^AMP are barely converted to m^6^AD/TP, m^6,6^AD/TP, and i^6^AD/TP by AK (Fig. S5K). Furthermore, in *Adal* KO mice, only m^6^AMP, m^6,6^AMP, and i^6^AMP were accumulated at high levels, while their di- or triphosphate forms were undetectable (Fig. 5D). These results suggest that m^6^AMP, m^6,6^AMP, and i^6^AMP are the products of dead-end modified adenosine metabolism. Although our results cannot rule out the existence of trace amounts of m^6^AD/TP, m^6,6^AD/TP, and i^6^AD/TP, the extreme low abundance of these metabolites means they are unlikely to have a meaningful impact on physiological functions *in vivo*.

Collectively, our findings show that cells possess a sophisticated mechanism by which three specific modified adenosines generated from RNA catabolism are sequentially metabolized to IMP, and that dysregulation of this system can have significant pathological consequences.

### Limitations of the study

Although this study demonstrated that impaired metabolism of modified adenosines (m^6^A, m^6,6^A, and i^6^A) is the causative pathology of ADK deficiency, the therapeutic efficacy of this pathway remained unexplored due to technical limitations. Since systemic *Adk* KO mice exhibited early lethality (Fig. 2C), the therapeutic use of conditional KO mice should be the subject of future studies. Another limitation of this work is that we cannot exclude the possibility that other factors besides the cytotoxicity of modified adenosines may contribute to the impairment of lipid metabolism in ADK deficiency. The ligand activity of modified adenosines, such as m^6^A toward the adenosine A3 receptor, as we have reported previously^21^, remains to be further elucidated in the context of lipid metabolism.

Additionally, the human pathology caused by mutations in ADAL was not explored because this gene has not been reported. Interestingly, a recent study showed that ADAL is associated with a mixed ancestral structure in the regulatory region across multiple tissue types^70^. When we extracted missense mutations from a genomic database (gnomAD; https://gnomad.broadinstitute.org), 198 variants were identified. Among them, 193 (97.5%) are very rare coding variants (allele frequency <0.1%), and the allele frequencies of the remaining five mutations (D82G, R254Q, A95S, L302I, and N237S) are between 0.1% and 0.5%. None of the missense mutations are common, but further analysis in humans is needed to understand the impact of genetic variance on ADAL-related impaired glucose metabolism.

## Acknowledgements

We are grateful for suggestions and technical support from all members of the Department of Modomics Biology & Medicine, Tohoku University. We thank Y. Takahata and H. Miyamoto for technical assistance, and Natalie D. DeWitt for critical reading and language editing. This work was supported by funding provided by JSPS KAKENHI (Grant nos. 22H02813, 22H04628, and 23K18088 to A.O., 22H04922 (AdAMS) and 16H06276 (AdAMS) to K.A., 20H00422 and 23H04748 to S.S., 21H04758 to K.I., and 21H02659 and 21H05265 to F.-Y.W.), and AMED-CREST (21gm1410006h0001 to K.I.). This work was also supported by FOREST from JST (Grant nos. JPMJFR220K to A.O), ERATO JPMJER2002 from J.S.T. to F.Y.W.. Further support was obtained from the Takeda Science Foundation (to A.O. and F.Y.W.); the Uehara Memorial Foundation (to A.O.); the Sumitomo Foundation (to A.O.); the Naito Foundation (to A.O. and F.Y.W.); the Shiseido Female Researcher Science Grant (to A.O.; this grant was only used for cellular experiments); the Gout and uric acid foundation (to A.O.); and the Inamori foundation (to A.O.).

## Author contributions

The authors confirm their contribution to the paper as follows: Study conceptualization and design: A.O. and F.Y.W.; data collection: A.O., S.W, T.K., A.Y.-L.T., S.Y., K.A.; analysis and interpretation of results: A.O., S.W., A.Y.-L.T., K.A., S.S., K.I, and F.Y.W.; draft paper preparation: A.O., S.W, A.Y.-L.T., K.A., S.S., K.I, and F.Y.W.; project supervision: A.O., K.I., and F.Y.W. All authors reviewed the results and approved the final version of the paper.

## STAR★Methods

### Key Resource Table

| REAGENT or RESOURCE | SOURCE | IDENTIFIER |
| --- | --- | --- |
| Antibodies |  |  |
| Phospho-Acetyl-CoA Carboxylase (Ser79) (D7D11) | Cell Signaling Technology | Cat#11818S<br>RRID:AB_2687505 |
| Acetyl-CoA Carboxylase (C83B10) | Cell Signaling Technology | Cat#3676S<br>RRID:AB_2219397 |
| AMPK alpha (phospho Thr172) | GeneTex | Cat#GTX52341<br>RRID:AB_2887793 |
| AMPKα (D5A2) | Cell Signaling Technology | Cat#5831S<br>AB_10622186 |
| Fatty Acid Synthase | Abcam | Cat#ab22759<br>RRID:AB_732316 |
| SCD1 | Cell Signaling Technology | Cat#2438S<br>RRID:AB_823634 |
| SREBP-1 (A-4) | Santa Cruz Biotechnology | Cat#sc-365513<br>RRID:AB_10843812 |
| CPT1A [8F6AE9] | Abcam | Cat#ab128568<br>RRID:AB_11141632 |
| CPT2 [EPR13626] | Abcam | Cat#ab181114<br>RRID:AB_2687503 |
| LC3B | Cell Signaling Technology | Cat#2775S<br>RRID:AB_915950 |
| SQSTM1/p62 (D6M5X) | Cell Signaling Technology | Cat#23214S<br>RRID:AB_2798858 |
| β-actin | MBL International | Cat#M177-3<br>RRID:AB_10697039 |
| Adenosine kinase | Thermo Fisher Scientific | Cat#PA5-27399<br>RRID:AB_2544875 |
| ADAL | Proteintech | Cat# 27369-1-AP<br>RRID:AB_2880855 |
| LC3B [EPR18709] | Abcam | Cat#ab225383<br>RRID:AB_2928102 |
| Bacterial and Virus Strains |  |  |
| One Shot TOP10 Chemically Competent *E. coli* | Thermo Fisher Scientific | Cat#C404003 |
| NEB 10-beta Competent *E. coli* | New England BioLabs | Cat#C3019 |
| Chemicals, Peptides, and Recombinant Proteins |  |  |
| *N*<sup>6</sup>-Threonylcarbamoyladenosine | Medicilon | Custom-synthesized |
| 2-Methylthio-*N*<sup>6</sup>-threonylcarbamoyladenosine | Medicilon | Custom-synthesized |
| 2-Methylthio-*N*<sup>6</sup>-isopentenyladenosine | Medicilon | Custom-synthesized |
| *N*<sup>6</sup>-2'-O-Dimethyladenosine | Toronto Research Chemicals | Cat#D447415;<br>CAS:57817-83-1 |
| 2'-O-Methyladenosine | Tokyo Chemical Industry | Cat#M2291;<br>CAS:2140-79-6 |
| 1-Methyladenosine | Cayman Chemical | Cat#16937;<br>CAS:15763-06-1 |
| 2'-O-Methylinosine | Santa Cruz Biotechnology | Cat#sc-283498;<br>CAS:3881-21-8 |
| 1-Methylinosine | Santa Cruz Biotechnology | Cat#sc-483758;<br>CAS:2140-73-0 |
| *N*<sup>6</sup>-Methyladenosine | Abcam | Cat#ab145715;<br>CAS:1867-73-8 |
| *N*<sup>6</sup>,*N*<sup>6</sup>-dimethyladenosine | MedChem Express | Cat#HY-101984;<br>CAS:2620-62-4 |
| *N*<sup>6</sup>-Isopentenyladenosine | Toronto Research Chemicals | Cat#I821840;<br>CAS: 7724-76-7 |
| *N*<sup>4</sup>-Acetylcytidine | Combi-Blocks | Cat#QB-9019;<br>CAS:3768-18-1 |
| 2'-O-Methylcytidine | Tokyo Chemical Industry | Cat#M2317;<br>CAS:2140-72-9 |
| 5-Methylcytidine | Tokyo Chemical Industry | Cat#M1931;<br>CAS:2140-61-6 |
| 5-Formylcytidine | Toronto Research Chemicals | Cat#F698965;<br>CAS:148608-53-1 |
| 3-Methylcytidine (methosulfate) | Cayman Chemical | Cat#21064;<br>CAS: 21028-20-6 |
| 2'-O-Methyl-5-methylcytidine | Granlen | Cat#100726<br>CAS: 113886-70-7 |
| 5-(Hydroxymethyl)cytidine | Tokyo Chemical Industry | Cat#H1863;<br>CAS RN: 19235-17-7 |
| *N*<sup>2</sup>,*N*<sup>2</sup>-Dimethylguanosine | MedChem Express | Cat#HY-113137;<br>CAS:2140-67-2 |
| *N*<sup>2</sup>-Methylguanosine | Biosynth Carbosynth | Cat#NM35522;<br>CAS:2140-77-4 |
| 1-Methylguanosine | Santa Cruz Biotechnology | Cat#sc-500889;<br>CAS:2140-65-0 |
| 2'-O-Methylguanosine | Tokyo Chemical Industry | Cat#M2318;<br>CAS:2140-71-8 |
| 7-Methylguanosine | Sigma-Aldrich | Cat#M0627;<br>CAS:20244-86-4 |
| Pseudouridine | Tokyo Chemical Industry | Cat#P2396;<br>CAS:1445-07-4 |
| N1-Methylpseudouridine | MedChem Express | Cat#HY-112582;<br>CAS No.13860-38-3 |
| 3-(3-Amino-3-carboxypropyl) uridine | Alsachim | custom-synthesized |
| 5,6-Dihydrouridine | Toronto Research Chemicals | Cat#D449668;<br>CAS:5627-05-4 |
| 2'-O-Methyluridine | Tokyo Chemical Industry | Cat#M2290;<br>CAS:2140-76-3 |
| 3-Methyluridine | MedChem Express | Cat# HY-113138;<br>CAS:2140-69-4 |
| 5-Methyluridine | Tokyo Chemical Industry | Cat#M1405;<br>CAS: 1463-10-1 |
| 5-Taurinomethyluridine | Alsachim | custom-synthesized |
| 2-Thiouridine | Toronto Research Chemicals | Cat#T384000;<br>CAS: 20235-78-3 |
| 5-Taurinomethyl-2-thiouridine | Alsachim | custom-synthesized |
| Adenosine monophosphate | MedChem Express | Cat#HY-A0181<br>CAS: 61-19-8 |
| N6-Methyladenosine 5'-monophosphate sodium salt | Sigma-Aldrich | Cat#M2780<br>CAS:81921-35-9 |
| N6-Dimethyladenosine-5'-monophosphate sodium salt | Tokyo Chemical Industry | custom-synthesized |
| N6-Isopentenyladenosine-5'-monophosphate sodium salt | Biosynth | Cat#NI30240<br>CAS: 1402800-76-3 |
| Inosine 5'-monophosphate disodium salt hydrate | Tokyo Chemical Industry | Cat#I0036<br>CAS: 352195-40-5 |
| Uridine | Sigma-Aldrich | Cat#U3750;<br>CAS:58-96-8 |
| Cytidine | Sigma-Aldrich | Cat#C122106;<br>CAS:65-46-3 |
| Guanosine | Sigma-Aldrich | Cat#G6752;<br>CAS: 118-00-3 |
| Adenosine | Sigma-Aldrich | Cat#A9251;<br>CAS:58-61-7 |
| Inosine | Sigma-Aldrich | Cat#I4125;<br>CAS:58-63-9 |
| Hydrogen Peroxide | FUJIFILM Wako Pure Chemical | Cat#081-04215 |
| Mitomycin C | Nacalai Tesque | Cat#20898-21 |
| Staurosporine | FUJIFILM Wako Pure Chemical | Cat#197-10251 |
| Methotrexate | FUJIFILM Wako Pure Chemical | Cat#135-13573 |
| Deoxycoformycin | Sigma-Aldrich | Cat#116860 |
| ABT-702 dihydrochloride | MedChem Express | Cat#HY-103161 |
| Opti-MEM | Thermo Fisher Scientific | Cat#31985-070 |
| Polyethylenimine (PEI) Max, (Mw 40,000) | Polysciences | Cat#24765-1 |
| Recombinant Human Adenosine Deaminase | R&D Systems | Cat#7048-AD |
| Recombinant Human Adenosine Kinase | R&D Systems | Cat#8024-AK |
| Recombinant Human Adenylate Kinase 1 Protein | Novus Biologicals | Cat#NBP1-50855 |
| TALON Metal Affinity Resin | Clontech | Cat#635501 |
| BODIPY 493/503 | Invitrogen | Cat#D3922 |
| Critical Commercial Assays |  |  |
| Pierce BCA Protein Assay Kit | Thermo Fisher Scientific | Cat#23225 |
| PrimeScript RT-PCR kit | TAKARA | Cat#RR014A |
| SYBR Premix EX Taq II | TAKARA | Cat#RR820A |
| KOD-Plus-Mutagenesis kit | TOYOBO | Cat#SMK-101 |
| SapphireAmp Fast PCR Master Mix | TAKARA | Cat#RR350A |
| Deposited Data |  |  |
| N/A | N/A | N/A |
| Experimental Models: Cell Lines |  |  |
| HEK293A | Inoue et al., 2012 | N/A |
| HEK293T | ATCC | N/A |
| HEK293FT | ATCC | N/A |
| HepG2 | ATCC | N/A |
| MCF7 | JCRB | JCRB0134 |
| THP1 | JCRB | JCRB0112.1 |
| U87 | ATCC | N/A |
| MDA231 | JCRB | JCRB1559 |
| HConF | Sciencell | SCR 6570 |
| MEF | BRC | RCB2710 |
| Experimental Models: Organisms/Strains |  |  |
| C57BL/6J male mice | Japan Clea | N/A |
| Oligonucleotides |  |  |
| siRNA targeting: human Perilipin 1 (PLIN1) | Thermo Fisher Scientific | siRNA ID: s10661 |
| siRNA targeting: human Perilipin 2 (PLIN2) | Thermo Fisher Scientific | siRNA ID: s1056 |
| siRNA targeting: human Perilipin 3 (PLIN3) | Thermo Fisher Scientific | siRNA ID: s19954 |
| siRNA targeting: human Perilipin 4 (PLIN4) | Thermo Fisher Scientific | siRNA ID: s59654 |
| Software and Algorithms |  |  |
| Prism 10.0 | GraphPad software | https://www.graphpad.com/<br>RRID:SCR_002798 |
| Other |  |  |
| Amicon Ultra-0.5 Centrifugal Filter Unit | Merck | Cat#UFC500396 |
| SpeedVac Vacuum Concentrators | Thermo Fisher Scientific | Cat#SPD1010 |
| Inertsil ODS-3 Column | GL Science | Cat#5020-84655 |
| Real Time PCR System | Qiagen | Rotor-Gene Q |

### RESOURCE AVAILABILITY

#### Lead contact

Further information and requests for reagents and resources should be directed to, and will be fulfilled by, the corresponding authors Akiko Ogawa or Fan-Yan Wei.

#### Materials availability

All plasmids, cell lines, and reagents generated in this study are available from the corresponding authors via a completed Materials Transfer Agreement.

#### Data availability

The RNA-seq data (raw data and processed data) are available in the GEO repository (accession number: GSE262450).

### EXPERIMENTAL MODEL AND STUDY PARTICIPANT DETAILS

#### Cell lines

All cells were cultured at 37°C in an atmosphere of 5% CO2. HEK293A, HEK293T, and HEK293FT cells were cultured in Dulbecco’s modified Eagle’s medium (DMEM; Wako, Osaka, Japan) supplemented with 10% heat-inactivated fetal bovine serum (FBS; BioWest, Nuaillé, France). Transient transfections were performed using polyethylenimine (PEI) transfection reagent (Polysciences, Pennsylvania, USA). Transfection efficiency was analyzed by qPCR.

## METHOD DETAILS

### Sample preparation for mass spectrometry analysis

Samples for measuring metabolites including modified and unmodified nucleosides were prepared for mass spectrometry analysis as previously described^71^. Briefly, metabolites in extracellular fluids of cultured cells were extracted with ice-cold methanol. The suspension was then centrifuged at 16,000 g for 10 min at 4°C. After transfer of the top layer, ultrapure water was added and the suspension was ultrafiltered using an Amicon Ultra-0.5 3 kDa cut-off ultrafiltration tube (Merck Millipore, Massachusetts, USA). The ultrafiltrate was concentrated using a SpeedVac vacuum concentrator (Thermo Fisher Scientific, Massachusetts, USA), and the concentrated filtrate was dissolved in ultrapure water and subjected to LCMS analysis.

Intracellular metabolites in cultured cells were extracted with ice-cold methanol. Pre-chilled chloroform and ultrapure water was sequentially added to the suspension, and centrifuged at 10,000 g for 5 min at 4°C. The top layer was ultrafiltered using an ultrafiltration tube. The ultrafiltrate was concentrated using a vacuum concentrator, and the concentrated filtrate was dissolved in ultrapure water and subjected to LC-MS analysis.

For mouse experiments, C57BL/6J male mice were purchased from Japan clea (Tokyo, Japan). All animal studies were approved by the Institutional Laboratory Animal Care and Use Committee of Tohoku University (authorization no. 2019-047-01). Blood was collected into EDTA coated tubes on ice and centrifuged at 300 × g for 20 minutes to separate plasma, that was stored in aliquots at −80°C until analysis. Urine was immediately frozen at −80°C until further analysis. Metabolites in extracellular fluids of biological samples were extracted with ice-cold methanol/chloroform/ultrapure water. The suspension was then centrifuged at 16,000 × g for 3 min at 4°C. After transfer of the top layer, ultrapure water was added and the suspension was centrifuged at 16,000 g for 3 min at 4°C. After centrifugation, the supernatant was ultra-filtered using an ultrafiltration tube. Tissues were homogenized in pre-chilled buffer containing an equal amount of water and methanol using TissueRuptor II with a disposable probe (Qiagen, Hilden, Germany). The homogenate was centrifuged at 16,000 × g for 15 min at 4°C, and the supernatant was transferred to an ultrafiltration tube. The ultrafiltrate was concentrated using a vacuum concentrator, and the concentrated filtrate was dissolved in ultrapure water and subjected to LC-MS analysis.

### Mass spectrometry

Quantitative modified nucleoside/nucleotide analysis was performed using an Orbitrap Exploris 240 mass spectrometer (Thermo Fisher Scientific), a Q Exactive instrument (Thermo Fisher Scientific), or an LCMS-8060NX instrument (Shimadzu, Kyoto, Japan) as previously described^71^ with slight modifications. Samples were injected into an Inertsil ODS-3 column (GL Science, Tokyo, Japan), a Discovery HS F5 HPLC column (Sigma, MO, USA), or an Intrada Organic Acid column (Imtakt, Kyoto, Japan) according to the type of nucleosides/nucleotides. The mobile phase consisted of 5 mM ammonium acetate pH 5.3 in water and 100% acetonitrile. Calibration curves using various concentrations of the nucleosides to be measured were obtained in each analytical run.

For non-targeted metabolomics, a Discovery HS F5 HPLC column (Sigma) was used to analyze cation metabolites, and a seQuant ZIC-HILIC column was used to analyze anion metabolites, both according to the manufacturer’s instructions. Orbitrap Exploris 240 was set to full mass scan mode at a resolution of 120,000 in combination with data-dependent analysis using ms/ms mode. Data analysis was performed using Compound Discoverer 3 (Thermo Fisher Scientific) following standard procedures. Principal component analysis was performed using the MetaboAnalyst platform (https://www.metaboanalyst.ca)^72^.

### Evaluation of cell viability

Cell viability was assessed by WST-8 assay using a Cell Counting Kit-8 (CCK-8; Dojindo, Kumamoto, Japan) according to the manufacturer’s protocol. After treatment with each nucleoside, WST-8 solution was added to each well, and plates were incubated for 1 h. The absorbance at 450 nm was then measured using a SpectraMax ABS microplate reader (Molecular Devices, CA, USA).

### Evaluation of cytotoxicity

Cell viability was assessed using an LDH assay kit (Dojindo) according to the manufacturer’s protocol. After treatment with each nucleoside, working solution was added to each well, and plates were incubated for 30 min at room temperature under shaded light. Stop solution was added to each well, and the absorbance at 490 nm was measured using a SpectraMax ABS microplate reader (Molecular Devices).

### Compound screening for metabolic regulation of m^6^A, m^6,6^A, and i^6^A

Metabolic regulation of m^6^A, m^6,6^A, and i^6^A was screened as described below. HEK293A cells were seeded into 96-well plates. After 24 h, culture medium was replaced with fresh medium containing each compound from the Kinase Screening Library (#10505; Cayman, Michigan, USA) and the Cellular Metabolism Screening Library (#33705; Cayman) at the desired concentration. After 1 h, a mixture of m^6^A, m^6,6^A, and i^6^A was added (final concentration 1 µM). The supernatant was collected after 12 h and subjected to LC-MS analysis to determine the stability of m^6^A, m^6,6^A, and i^6^A.

### Generation of labeled nucleosides and clearance analysis

To label RNA modifications with stable isotope in cells, HEK293A cells were cultured for 7 generations in D-glucose (U-^13^C6)-containing medium. Total RNA was purified and digested with nuclease P1 (Wako) and bacterial alkaline phosphatase (BAP; TAKARA, Shiga, Japan). The resulting modified nucleosides were exogenously added to cells. The modified nucleosides contained ^13^C-labeled ribose, which is derived from D-glucose (U-^13^C6). Decay of the labeled modified nucleosides was measured.

### Generation of knockout (KO) cells by CRISPR-Cas9

Single-guide RNAs (sgRNAs) were designed to target the human ADK or ADAL genomic locus. The sgRNA sequence driven by the U6 promoter was cloned into a lentiCRISPRv2 vector that also expresses Cas9 as previously described^73^. Lentiviral particles were produced in HEK293FT cells by co-transfection of the targeting vector with vectors expressing Gag, Pol, Rev, Tat, and VSV-G genes. The lentiviral plasmid DNA was then packed into lentivirus for infection of HEK293A cells. Infected cells were selected in puromycin for 2 weeks before single colonies were chosen and tested by DNA sequencing, qPCR, and western blotting.

### Generation of KO mice by CRISPR-Cas9

*Adk* KO and *Adal* KO mice were generated using CRISPR-Cas9 following standard procedures at the Institute of Resource Development and Analysis, Kumamoto University. The crRNAs with Cas9 recombinant protein were used to delete genomic regions including exons 4−6 of the mouse Adk gene, and exons 3−10 of the mouse Adal genes. All animals were housed in the animal facility of the Institute of Development, Aging and Cancer, Tohoku University (Miyagi, Japan). The Animal Ethics Committee of Tohoku University reviewed and approved all animal procedures (Approval ID: 2021-007-03). Animals were housed at 25°C under a 12 h light and 12 h dark cycle. The sequences of crRNAs used to generate *Adk* and *Adal* KO mice are as follows, with F0 mice backcrossed with C57BL6 mice for at least three generations: crRNA 1 for Adal gene: 5’-TCAGCTTACTACTAGTGCTG crRNA 2 for Adal gene: 5’-TCAGCTTACTACTAGTGCTG crRNA 1 for Adk gene: 5’-CACAAACATGAACATAACTG crRNA 2 for Adk gene: 5’-AAAAGCAGTAGACCCGAAGG

### Expression and purification of recombinant proteins

AG1 competent cells were transformed with the pCA24N-based plasmids encoding ADK or ADAL with each mutation, or truncated ACC comprising the first 100 amino acids which contains the phosphorylation site targeted by AMPK. The transformed cells were grown at 30°C in 200 mL of LB medium. When the optical density at 660 nm (OD660) reached 0.7, IPTG was added to the culture medium to a final concentration 1 mM to induce gene expression. After cultivation overnight at 25°C, cells were harvested, resuspended in 40 mL of lysis buffer (20 mM imidazole, 0.3 M NaCl, 50 mM sodium phosphate, supplemented with protease inhibitor cocktail; Nacalai Tesque, Kyoto, Japan) and lysed on ice using a Multi-beads Shocker (Yasui Kikai, Osaka, Japan). After centrifugation, the soluble fraction of the supernatant was purified using TALON metal affinity resin (TAKARA) following the manufacturer’s instructions, and proteins were eluted with buffer containing 250 mM imidazole, 0.3 M NaCl, and 50 mM sodium phosphate. Protein concentrations were determined using a BCA Protein Assay Kit (Thermo Fisher Scientific) with bovine serum albumin as the standard protein.

### *In vitro* phosphorylation assay

For ADK assay, purified recombinant human ADK with each mutation (5 ng/µL) was incubated for 10 min at 37°C in buffer containing ATP (1 mM), 10× assay buffer (250 mM HEPES, 1.5 M NaCl, 100 mM MgCl2, and 100 mM CaCl2), and varying concentrations of each substrate. Reactions were acidified by adding glacial acetic acid, and products were detected by LC-MS analysis. For AMPK assay, recombinant AMPK (α1, β1, and γ1, 0.5 ng/µL, Sigma), purified truncated ACC (250 ng/µL), ATP (50 µM), AMP (1 µM), and varying concentrations of modified AMPs were incubated for 15 min at 37°C. SDS-PAGE sample buffer (5×) was added to each product and samples were analyzed by SDS-PAGE as described below.

### Molecular docking

All molecular docking simulations in this study were performed with AutoDock Vina^74^. For docking of modified adenosine, the crystal structure of human ADK complexed with adenosine (PDB ID: 1BX4) was used as the target molecule. For docking of modified AMPs with ADAL, the target molecule was a homology model of human ADAL built with Swiss-model^75^ based on the crystal structure of m^6^AMP-bound AtADAL (PDB ID: 6ijn). For docking of modified AMPs with AMPK, the crystal structure of human AMPK in the active form (PDB ID: 4RER) was used as the target protein.

### Mutagenesis study

ADK and ADAL mutants were generated by introducing single point mutations into pCAG plasmids encoding human ADK and ADAL, using a KOD-Plus-Mutagenesis kit (TOYOBO, Tokyo, Japan), and confirmed by Sanger sequencing (Azenta, Tokyo, Japan). These plasmids were transfected into ADK KO or ADAL KO cells using polyethylenimine (PEI) agent.

### *In vitro* deamination assay

Purified recombinant human ADAL with each mutation (2.5 ng/µL) was incubated for 10 min at 37°C in buffer containing 10× assay buffer (250 mM HEPES, 1.5 M NaCl, 100 mM MgCl2, and 10 mM 2-mercaptoethanol) and varying concentrations of each substrate. Reactions were acidified by adding glacial acetic acid, and products were detected by LC-MS analysis.

### Generation of ADAL-deficient plants

The *mapda-1* (SALK_144851, Columbia-0 [Col-0] ecotype) and *mapda-2* (SALK_010573, Col-0 ecotype) tDNA insertion mutants were obtained from the Arabidopsis Biological Resource Center (ABRC; https://abrc.osu.edu)^76, 31^. The *mapda-3* (Pst00481, Nossen [No-0] ecotype) line was obtained from the Riken BioResource Research Center (BRC) Arabidopsis transposon-tagged lines collection (https://epd.brc.riken.jp/en/seed/transp).^77, 78^ Seeds were surface-sterilized, vernalized, and allowed to germinate on 1/2-strength Murashige & Skoog (MS) agar at 22°C under constant light for 7 days. Genomic DNA extraction and genotyping to detect the tDNA insertion were performed essentially as previously described^79^. PCR amplification was performed using SapphireAmp Fast PCR Master Mix (TAKARA) according to the manufacturer’s instructions. Primers used for genotyping are listed in Table S5. Plants homozygous or heterozygous for the *mapda* mutations, as well as the cognate wild-type (WT) plants (Col-0 and No-0), were grown on soil/vermiculite (1:1) mixture at 22°C under constant light for 1 month. Mature, healthy rosette leaves were flash-frozen in liquid nitrogen and homogenized for further analyses.

### Western blotting analysis

For blots of pACC, ACC, pAMPKα, and AMPKα, WT (HEK293A) and ADAL KO cells were washed twice with phosphate-buffered saline (PBS) and cultured in DMEM free from serum and amino acids, and whole-cell lysates were collected in freshly prepared RIPA lysis buffer (50 mM Tris-HCl pH 7.5, 150 mM NaCl, 1% NP-40, 0.5% sodium deoxycholate, 0.1% SDS, and 1 mM EDTA) containing protease inhibitors (Nacalai Tesque) and phosphatase inhibitors (Nacalai Tesque). For blots of FASN, SCD1, SREBP1, PPARα, CPT1a, CPT2, LC3-B, p62, ADK, ADAL, and β-actin, WT and ADK KO mice at postnatal day 5 or WT and ADAL KO mice at 20 weeks were sacrificed and proteins were extracted from liver tissues using RIPA lysis buffer containing protease inhibitors and phosphatase inhibitors. Lysates were sonicated and clarified by centrifugation. Protein concentrations were measured using a BCA Protein Assay Kit, adjusted to 1 mg/mL, and 30 μg of proteins were subjected to western blotting for each experiment. Samples were separated by SDS-PAGE and transferred onto polyvinylidene difluoride (PVDF) membranes. Blots were blocked with 5% non-fat milk in TBS-T (Tris-buffered saline with 0.1% Tween-20) for 1 h, then probed using various primary antibodies.

The primary antibodies and working dilutions used were as follows: pACC (AB_2687505), 1:1000; ACC (AB_2219397), 1:1000; pAMPKα (AB_2887793), 1:500; AMPKα (AB_10622186), 1:1000; FASN (AB_732316), 1:10000; SCD1 (AB_823634), 1:10000; SREBP1 (AB_10843812), 1:500; PPARα (AB_2885073), 1:500; CPT1a (AB_11141632), 1:1000; CPT2 (AB_2687503), 1:1000; LC3-B (AB_915950), 1:1000; p62 (AB_2798858), 1:1000; ADK (AB_2544875), 1:1000; ADAL (AB_2880855), 1:1000; β-actin (AB_10697039), 1:10000. Target antigens were incubated with appropriate horseradish peroxidase (HRP)-conjugated secondary antibodies (Cell Signaling, MA, USA) and visualized using ECL substrate (GE Healthcare, Illinois, USA). Phosphorylated protein levels were normalized against the total levels of the target protein. Levels of other protein levels were normalized against β-actin.

### Serum biochemistry

Biochemical parameters were analyzed using blood services (Oriental Yeast Co., Ltd., City, Country).

### Measurement of blood glucose

Mice were fasted for 14 h (8:00 pm to 10:00 am) followed by intraperitoneal injection of glucose (2 g/kg). Blood glucose was determined by an ACCU-CHEK glucometer (Roche).

### RNA-seq and gene ontology (GO)

Total RNA was isolated from WT and *Adk* KO mice at postnatal day 5 or WT and *Adal* KO mice at 32 weeks using TRIzol reagent (Thermo Fisher Scientific). RNA-seq was performed using an Illumina platform (Rhelixa, Tokyo, Japan). Raw data were analyzed using Galaxy (https://usegalaxy.org) as described elsewhere^80^. DESeq2 was used to analyze differences within two groups on the Galaxy platform. Differentially expressed genes (DEGs) were used to calculate the normalized enrichment score (NES) using Gene Set Enrichment Analysis (GSEA) following the instructions.

### Histology of mice

Sample tissues from liver were excised, fixed in 10% buffered neutral formalin (Wako) for 3 days, embedded in 2% paraffin, and sectionized. The samples were stained with hematoxylin and eosin (H&E), Elastica-Masson, and Oil Red O for histopathological examination under an APX100 microscope (Olympus, Tokyo, Japan).

### Lipid droplets

Lipid droplets were visualized using BODIPY staining (Thermo Fisher Scientific) as previously described^81^. In detail, HepG2 cells were seeded in 35 mm glass-bottomed cell culture dishes (Matsunami, Osaka, Japan). After 24 h of incubation, media was removed and replaced with fresh media containing vehicle, ABT702 (10 µM), or ABT702 (10 µM)+modified adenosine mixtures (100 nM of m^6^A, m^6,6^A, and i^6^A) for an additional 24 h incubation. Cells were then washed with PBS and stained with a solution containing 2 µM BODIPY for 15 min at 37°C. After washing with PBS, cells were fixed with 4% paraformaldehyde for 30 min. Cells were washed a further three times with PBS and stained with DAPI. Images were taken using a TCS-SP8 confocal microscope (Leica, Wetzlar, Germany).

## QUANTIFICATION AND STATISTICAL ANALYSIS

Statistical analyses were performed using GraphPad Prism 10 software (GraphPad Software, MA, USA) and methods are described in the legends of the figures. Symbols are mean values and error bars denote standard error of the mean (SEM) or standard deviation (SD). Representative results from at least two independent experiments are shown for every figure, unless stated otherwise in the figure legends. Additional details regarding the statistical analysis for each experiment are described in each figure legend.

## Supplementary Figures

**Figure S1.**
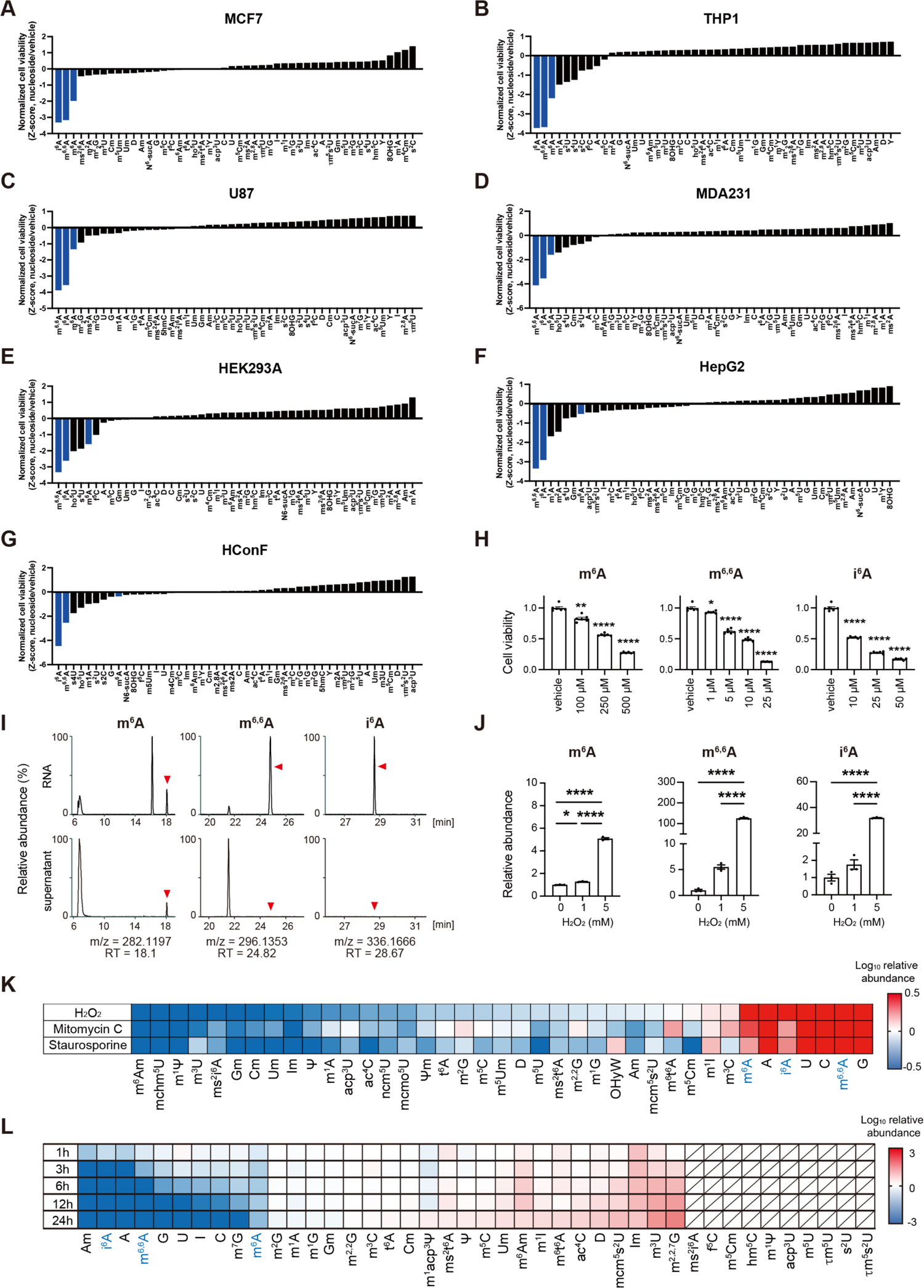
Dynamics and functions of modified nucleosides. Related to. Figure 1. (A−G) Normalized cell viability of human cell lines MCF7 (A), THP1 (B), U87 (C), MDA231 (D), HEK293A (E), HepG2 (F), and HConF (G) after 24 h incubation with each modified nucleoside (500 µM). n = 3−16. The sum of the results from the seven cell lines is shown in Figure 1A. Abbreviations for modified nucleosides are listed in Table S1. (H) Cell viability of MEF cells after 24 h incubation with different concentrations of modified nucleosides (n = 6). \**p* <0.05, \*\**p* <0.01, \*\*\*\**p* <0.0001, by one-way ANOVA followed by Dunnett’s test, compared with vehicle at the same incubation time. (I) Comparison of the abundance of m^6^A, m^6,6^A, and i^6^A in the supernatant vs. in RNA. Red arrows indicate the peak of each modified nucleoside. (J) Relative abundance of extracellularly released modified nucleosides in the supernatants of HEK293 cells after 24 h treatment with different concentrations of H_2_O_2_ compared to vehicle (n = 3). \**p* <0.05 and \*\*\*\**p* <0.0001, by one-way ANOVA followed by Tukey’s multiple comparison test. (K) Heatmap showing the relative abundance of extracellularly released modified nucleosides in the supernatants of HEK293 cells after 24 h treatment with different toxic compounds (5 mM H_2_O_2_, 0.1 mg/mL mitomycin C, 100 µM staurosporine) compared with vehicle. Averages of n = 3 are shown. (L) Heatmap showing the relative clearance of exogenously added labeled nucleosides from the supernatants of HEK293A cells after 24 h. Labeled nucleosides were obtained from enzymatic digestion of HEK293A total RNA passaged under D-glucose-^13^C_6_ for seven generations. Compared to 0 h. Averages of n = 3 are shown. Nucleosides with shaded areas are those that were not detected in this label system.

**Figure S2.**
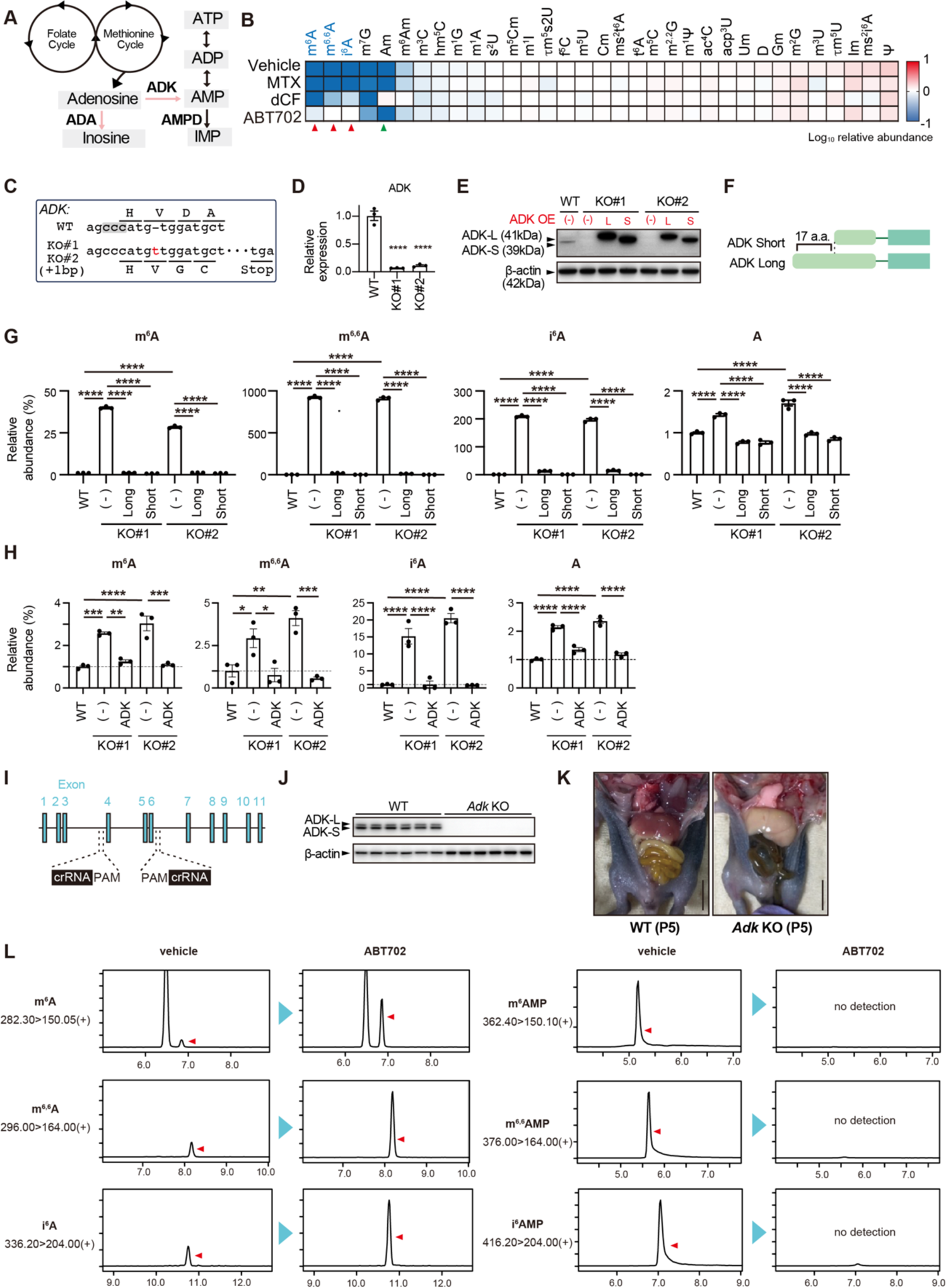
The three modified adenosines are substrates for adenosine kinase (ADK) but not adenosine deaminase (ADA). Related to Figure 2. (A) Intracellular adenosine metabolism. (B) Heatmap showing the relative clearance of exogenously added modified nucleosides (1 µM each) from the supernatants of HEK293 cells after 24 h, in the presence or absence of folate methotrexate (MTX, 100 µM), ADA inhibitor 2’-deoxycoformycin (dCF, 100 µM), or ADK inhibitor ABT702 (100 µM). Compared to 0 h. Averages of n = 3 are shown. Red arrows indicate that the rapid clearance of m^6^A, m^6,6^A, or i^6^A is reversed by ABT702. Green arrow indicates that the rapid clearance of Am is reversed by dCF. (C) Alignment of the genomic sequences of WT (HEK293A) and ADK KO cells edited by CRISPR-Cas9. The inserted nucleotide is highlighted in red. The PAM sequence is highlighted by a shaded box. Lines indicate reading frames. (D) Quantitative PCR (qPCR) detection of ADK expression in ADK KO cells. \*\*\*\**p* <0.0001, by one-way ANOVA followed by Dunnett’s test compared to WT. n = 3. Results are means ± SEM. (E) Protein levels of ADK examined by western blotting in WT, ADK KO, and ADK-reintroduced ADK KO cells for each isoform. (F) Isoforms of the human ADK gene. (G) Relative abundance of exogenously added extracellular m^6^A, m^6,6^A, and i^6^A (1 µM) in WT, ADK KO, and ADK-reintroduced ADK KO cells for each isoform. \*\*\*\**p* <0.0001, by one-way ANOVA followed by Tukey’s multiple comparison test. (H) Relative abundance of intracellular m^6^A, m^6,6^A, and i^6^A in WT, ADK KO, and ADK (Long)-reintroduced ADK KO cells. \**p* <0.05, \*\**p* <0.01, \*\*\**p* <0.001, and \*\*\*\**p* <0.0001, by one-way ANOVA followed by Tukey’s multiple comparison test. (I) Schematic view of the CRIPSR/Cas9 targeting strategy used for *Adk* KO mice. (J) Protein levels of ADK examined by western blotting in WT and *Adk* KO mice. n = 6, \*\*\*\**p* <0.0001 by unpaired t-test. (K) Representative photo of a WT (left) and an *Adk* KO (right) mouse at postnatal day 5. Scale bar = 5 mm. (L) Relative abundance of intracellular modified nucleosides and nucleotides 1 h after exogenous addition of m^6^A/m^6,6^A/i^6^A (1 µM) to HEK293A cells in the presence or absence of ABT702 (10 µM).

**Figure S3.**
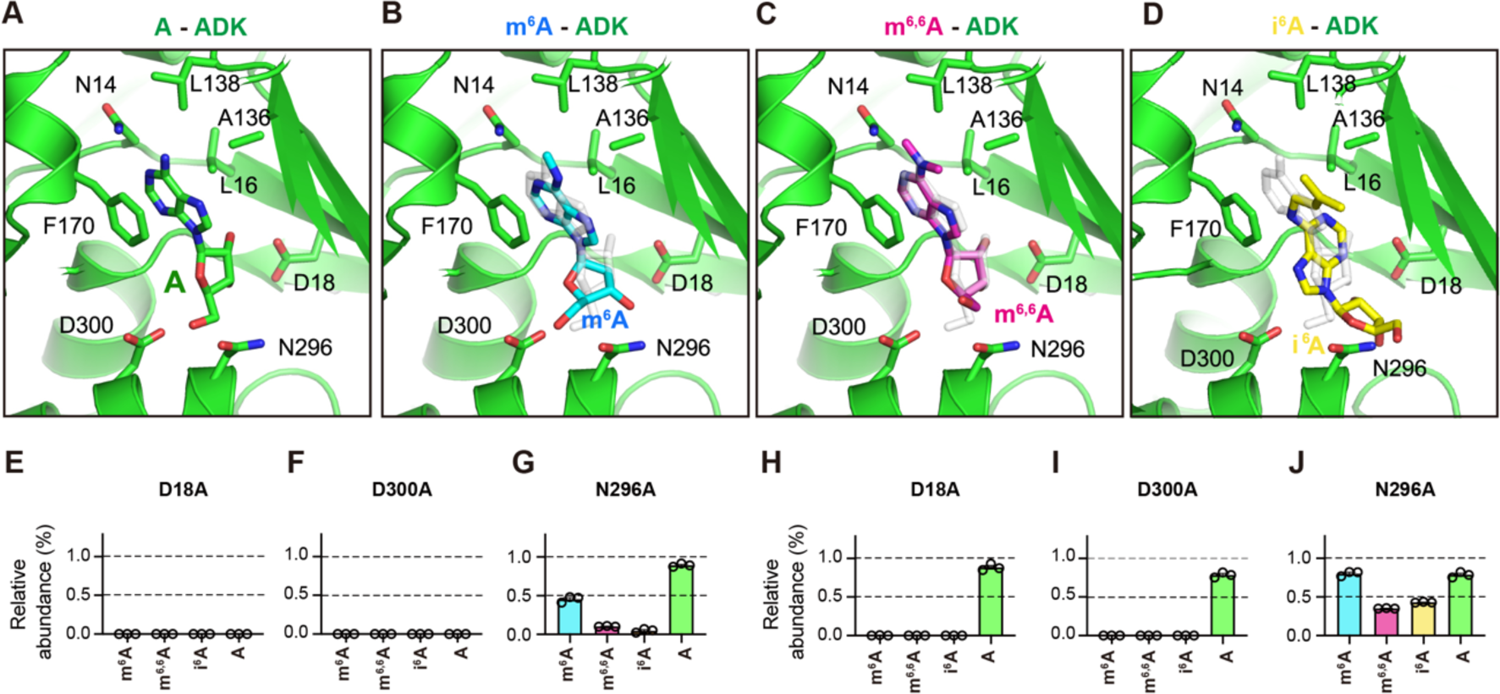
Molecular insights into of interaction between ADK and modified adenosines based on the crystal structure of the adenine bound ADK. Related to Figure 3. (A) Close-up view of the binding mode of adenosine in the catalytic site of ADK from the crystal structure of ADK in complex with adenosine (PDB: 1BX4). Oxygen and nitrogen atoms are colored in red and blue, respectively. (B-D) Close-up views of the predicted binding modes of (B) m^6^A (cyan), (C) m^6,6^A (magenta), and (D) i^6^A (yellow), bound to the catalytic site of ADK. The bound adenosine is also shown in light gray for comparison. (E−J) Relative abundance of products (i.e., m^6^AMP for m^6^A, m^6,6^AMP for m^6,6^A, i^6^AMP for i^6^A, and AMP for adenosine) from *in vitro* assays (E-G) or in cellular assays (H-J) with each ADK mutant (compared to the amount of each product generated by WT ADK). The x-axis shows the substrates used.

**Figure S4.**
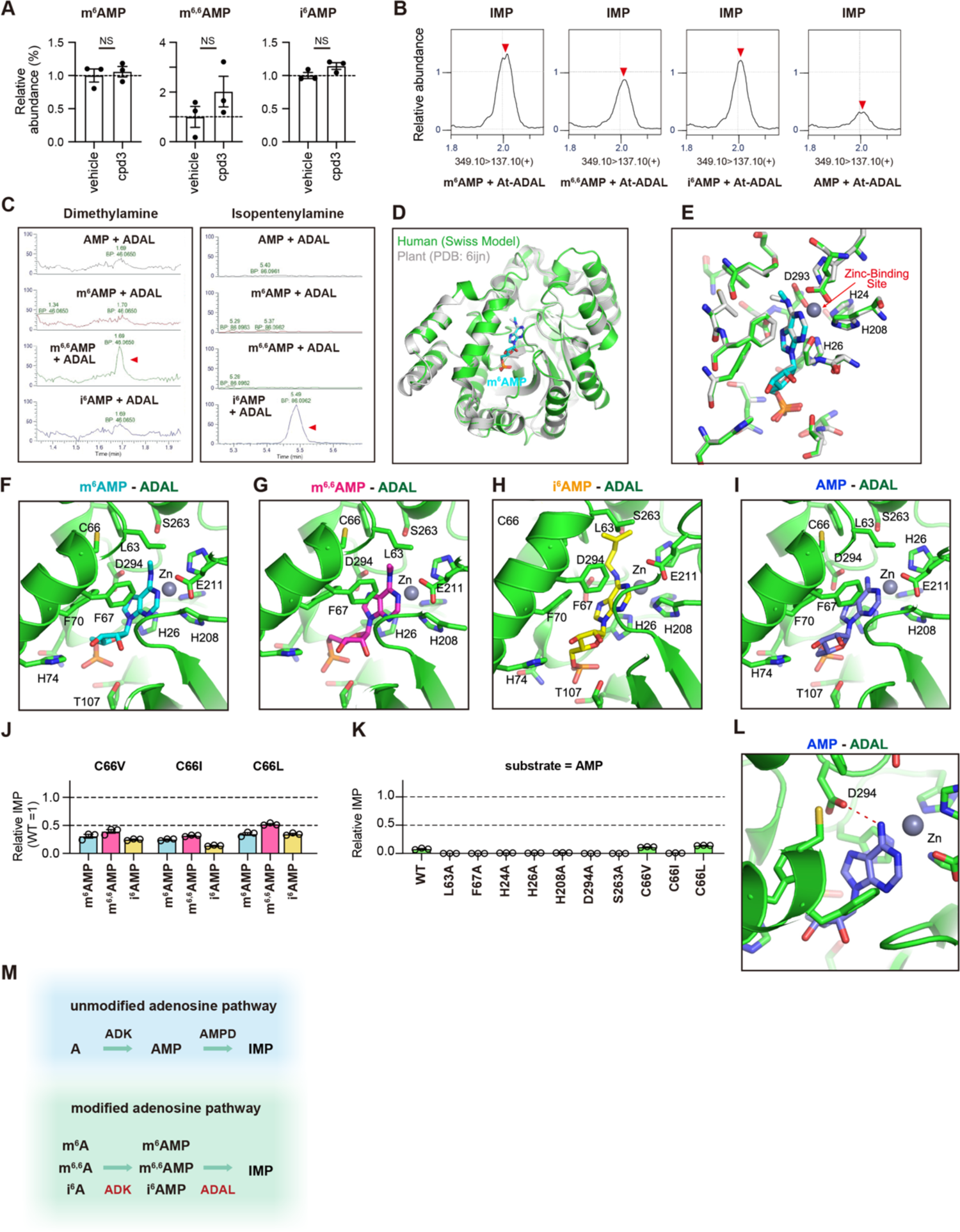
m^6^AMP, m^6,6^AMP, and i^6^AMP are substrates for adenosine deaminase-like (ADAL). Related to Figure 4. (A) Relative abundance of intracellular m^6^AMP, m^6,6^AMP, and i^6^AMP 24 h after exogenous addition of m^6^A, m^6,6^A, and i^6^A (1 µM each) to HEK293A cells in the presence or absence of Cpd3 (AMPD inhibitor, 10 µM). n = 3, unpaired t-test. (B) Relative abundance of IMP measured by LC-MS following *in vitro* deamination assay using At-ADAL protein. (C) Representative high-resolution mass spectra of dimethylamine (left) and isopentenylamine (right) following removal of the dimethyl moiety from m^6,6^AMP or the isopentenyl moiety from i^6^AMP by ADAL, respectively, detected using an Orbitrap in positive ion mode. (D) Homology modeling of human ADAL with m^6^AMP using SWISS MODEL (green) based on the plant ADAL structure complexed with m^6^AMP (gray, PDB: 6ijn). Oxygen, nitrogen, and phosphorus atoms are colored in blue, red, and orange, respectively. (E) Close up view of the bound m^6^AMP (cyan) with surrounding residues of the homology model of human ADAL (green), superimposed with those of plant ADAL complexed with m^6^AMP (light gray). (F-I) Close up views of the predicted binding mode of (F) m^6^AMP (cyan), (G) m^6,6^AMP (magenta), (H) i^6^AMP (yellow) and (I) AMP (slate blue) bound to human ADAL. (J) Relative abundance of IMP produced in *in vitro* deamination assays using ADAL C66V, C66I, and C66L mutants (compared to the amount of IMP generated by WT ADAL). The x-axis shows the substrates used. (K) Relative abundance of IMP produced in *in vitro* deamination assays using AMP and ADAL with each mutation (compared to the amount of IMP generated by WT ADAL and m^6^AMP as a substrate). (L) The enlarged view of predicted interactions in the docking simulation of the ADAL-AMP complex. Red dashed lines indicate hydrogen bonds between D294 and an amino group of AMP. (M) Pathways for unmodified and modified adenosines identified in this study. Unmodified adenosines are phosphorylated to AMP by ADK, and are deaminated to IMP by AMPD. Modified adenosines including m^6^A, m^6,6^A, and i^6^A are phosphorylated by ADK into m^6^AMP, m^6,6^AMP, and i^6^AMP, and further deaminated by ADAL into IMP.

**Fig. S5.**
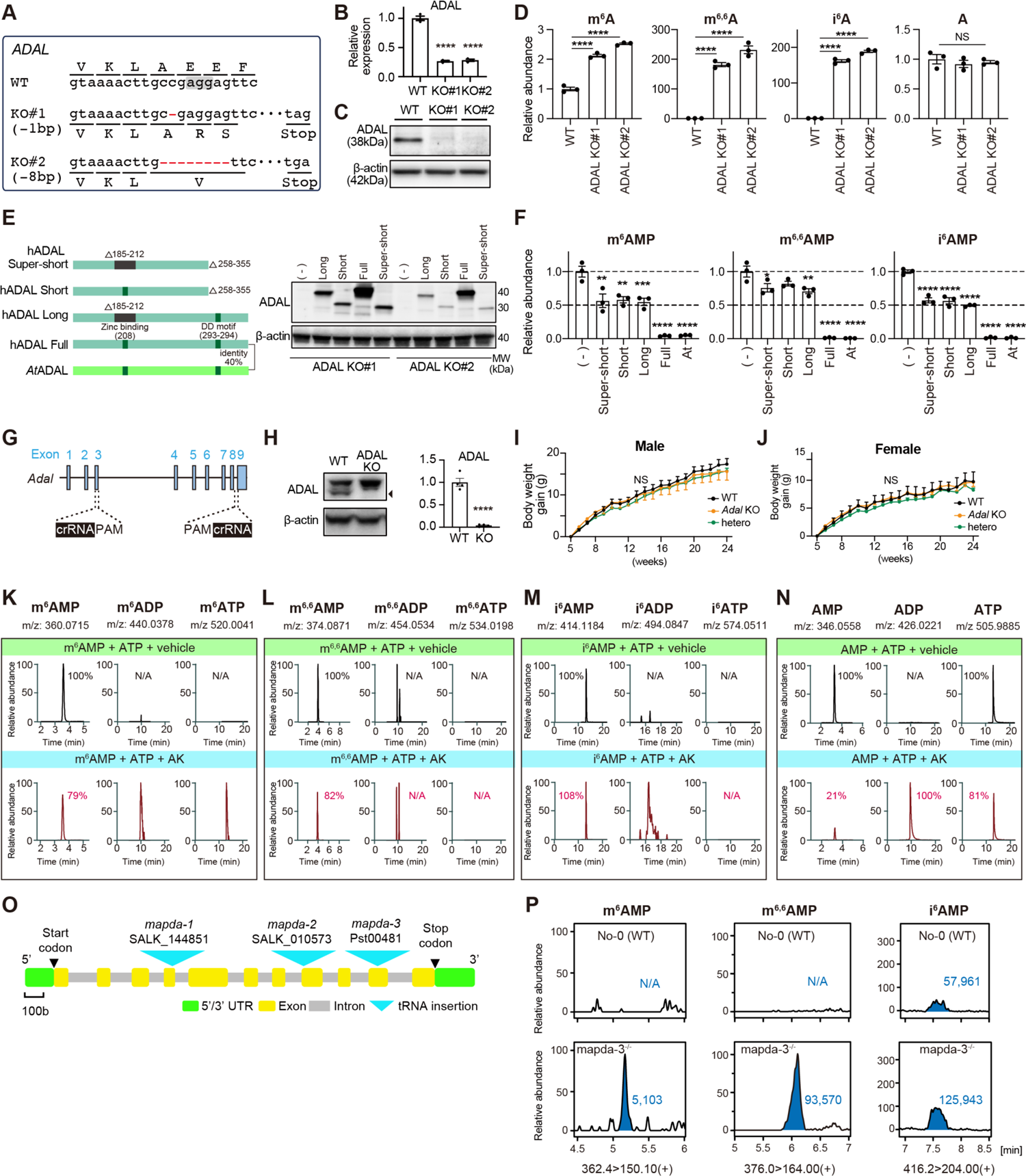
*In vivo* evidence of ADAL-mediated clearance of m^6^AMP, m^6,6^AMP, and i^6^AMP in *Arabidopsis thaliana* and mammals. Related to Figure 5. (A) Alignment of the genomic sequences of WT (HEK293A) and ADAL KO cells edited by CRISPR-Cas9. Deleted nucleotides are colored red. PAM sequences are highlighted by a shaded box. Lines indicate the reading frame. (B) qPCR detection of ADAL expression in ADAL KO cells. \*\*\*\**p* <0.0001, by one-way ANOVA followed by Dunnett’s test compared with WT. n = 3. Results are means ± SEM. (C) Protein levels of ADAL examined by western blotting in WT and ADAL KO cells. (D) Relative abundance of extracellular m^6^A, m^6,6^A, i^6^A, and adenosine in ADAL KO cells. \*\*\*\**p* <0.0001 by one-way ANOVA followed by Dunnett’s test compared with WT cells (n = 3). (E) (left) Isoforms of the human ADAL gene and *A. thaliana* (*At*-) ADAL. (right) Protein levels of ADAL examined by western blotting in WT, ADAL KO, and ADAL-reintroduced ADAL KO cells for each isoform. (F) Relative abundance of intracellular m^6^AMP, m^6,6^AMP, and i^6^AMP in ADAL KO cells (clone#2) and ADAL-reintroduced ADAL KO cells by each human isoform or At-ADAL. \*\**p* <0.01, \*\*\**p* <0.001, and \*\*\*\**p* <0.0001, by one-way ANOVA followed by Dunnett’s test compared with ADAL KO without reintroduction. (G) Schematic view of the CRIPSR/Cas9 targeting strategy used for *ADAL* KO mice. (H) Protein levels of ADAL examined by western blotting in WT and *ADAL KO* mice (n = 4, \*\*\*\**p* <0.0001 by unpaired t-test). (I-J) Body weight change of male (I) and female (J) WT (black), *Adal*^+/-^ (hetero, green), and *Adal* KO (orange) mice. Two-way repeated measure ANOVA was performed. (K−N) Levels of modified AMPs, ADPs, and ATPs in *in vitro* assays using human adenylate kinase (AK) with m^6^AMP (K), m^6,6^AMP (L), i^6^AMP (M), and AMP (N) as substrates. (O) Exon structure of the Arabidopsis *mapda* (At4g04880, an Arabidopsis ADAL orthologue) locus, and the tDNA insertion positions of the *mapda* mutants used in this study. (P) Representative abundance of modified AMPs obtained from rosette leaf extracts of 1-month-old WT (Nossen) and mapda-3^-/-^ Arabidopsis plants. n = 3.

**Figure S6.**
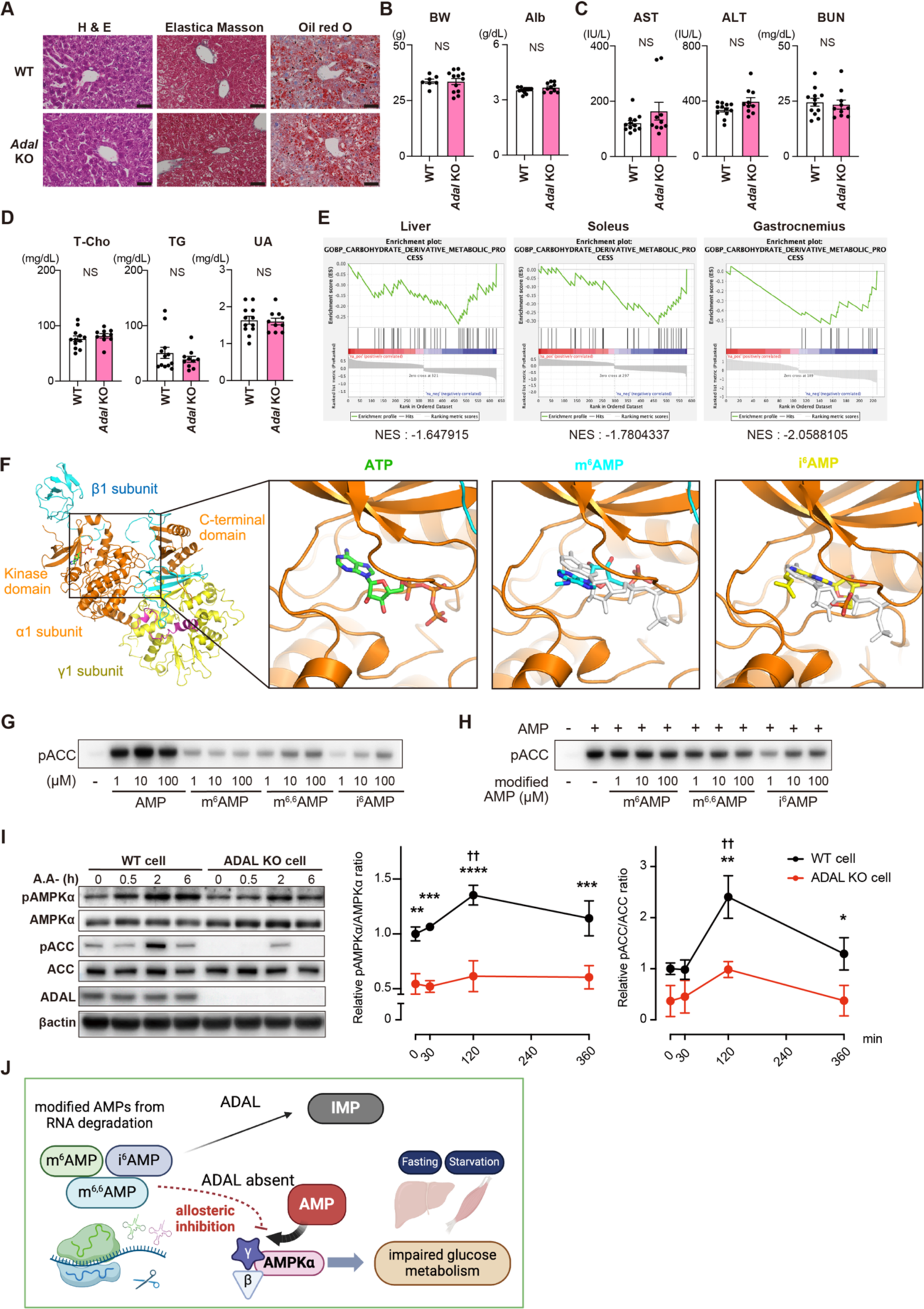
Carbohydrate metabolism is mildly impaired in *Adal* KO mice. Related to Figure 6. (A) H&E staining, Elastica Masson staining, and Oil red O staining of liver in WT and *Adal* KO mice. Scale bar = 50 µm. (B) Body weight and serum albumin of male WT (n = 7) and *Adal* KO (n = 12) littermates at 30 weeks of age. Results are means ± SEM. Unpaired t-test. (C-D) Blood test results of male WT (n = 12) and *Adal* KO (n = 10) littermates at 30 weeks of age. Results are means ± SEM. \**p* <0.05 between WT and *Adal* KO mice, by unpaired t-test. (F) Gene set enrichment analysis (GSEA) of each organ of *Adal* KO vs. WT mice. RNA-Seq results show negative enrichment of gene sets involved in carbohydrate derivative metabolic processes. (G) Docking of the modified AMPs with the kinase domain of the active AMPK. (Left) The crystal structure of active AMPK with a modeled ATP molecule, based on the ATP bound structure of cAMP-dependent protein kinase (PDB, 1ATP), which is similar to the kinase domain of AMPK. The C-terminal and kinase domains of the α1 subunit are in orange, the β1 subunit is in light blue, and the γ1 subunit is in yellow. (Right) Enlarged views of the ATP-binding site in the kinase domain. ATP is shown in green, m^6^AMP in cyan, and i^6^AMP in yellow. ATP is shown in light gray for comparison in the models of m^6^AMP and i^6^AMP-bound states. (G, H) SDS-PAGE of phosphorylated acetyl-CoA carboxylase 1 (ACC) following *in vitro* phosphorylation assay to determine AMPK activity. (G) Representative western blot for incubation of ACC, ATP, recombinant AMPK, and AMP/m^6^AMP/m^6,6^AMP/i^6^AMP. (H) Representative western blot for the incubation of ACC, ATP, recombinant AMPK (α1/β1/γ1), and AMP plus m^6^AMP/m^6,6^AMP/i^6^AMP. (I) Representative western blot in the time course analysis of amino acid starvation in WT and ADAL KO cells (n = 3). Densitometry is shown for relative pAMPKα/AMPKα and pACC/ACC (right). \**p* <0.05, \*\**p* <0.01, \*\*\**p*<0.001, \*\*\*\**p* <0.0001 vs. WT, by two-way ANOVA followed by uncorrected Fisher’s LSD. Results are means ± SEM. (J) Schematic diagram of the mechanism by which modified AMPs derived from RNA degradation allosterically inhibit AMPK activation, resulting in impaired glucose metabolism.

**Figure S7.**
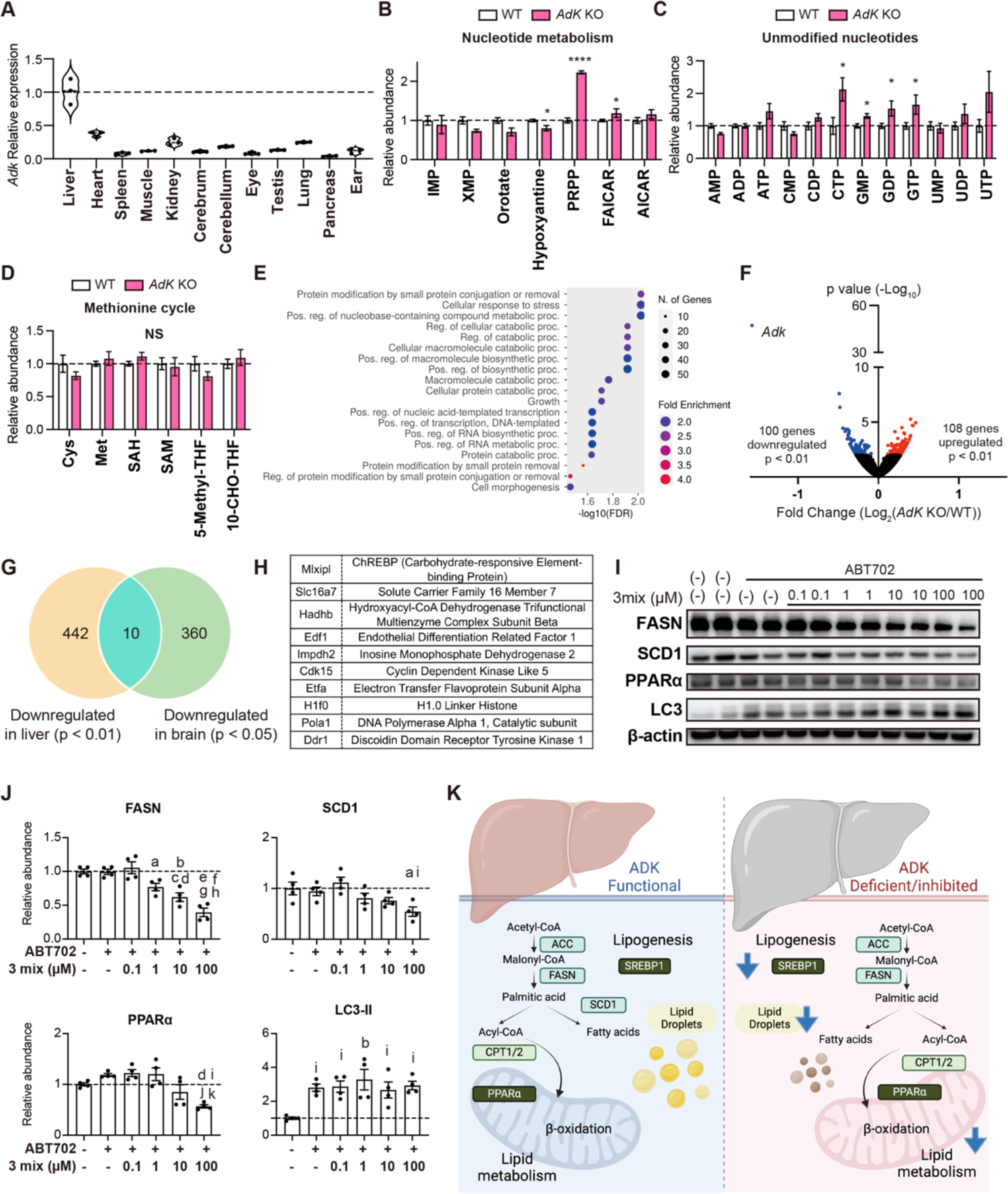
ADK deficiency disrupts lipid metabolism. Related to Figure 7. (A) Relative gene expression profile of *Adk* in the organs of mice determined by qPCR (n = 3). 18S rRNA was used for normalization of PCR products. (B−D) Metabolomics results showing (B) nucleotide metabolism, (C) unmodified nucleotides, and (D) the methionine cycle of liver from WT (n = 7) and *Adk* KO (n = 3) mice sacrificed at postnatal day 5. \**p* <0.05 and \*\*\*\**p* <0.0001, between WT and *AdK* KO mice by unpaired t-test. Results are means ± SEM. (E) The top 20 enriched pathways of upregulated genes identified using GO. Dot size represents the number of enriched genes and dot color represents the fold enrichment. (F) Volcano plot of RNA-seq results comparing gene expression in the brain of *Adk* KO and WT mice (>2-fold change, adjusted *p* <0.01 by DESeq2). (G, H) Venn diagram (G) and table (H) showing the overlap of downregulated genes in liver (*p* <0.01) and brain (*p* <0.05). (I, J) Representative blots (I) of the levels of proteins related to lipid metabolism or autophagy in HepG2 cells. HepG2 cells were incubated with ABT702 (10 µM) for 1 h, followed by the addition of the indicated concentrations of the three modified adenosine mixtures (m^6^A, m^6,6^A, and i^6^A) for 24 h (n = 4). (J) Densitometry results. a, *p* <0.01 vs. ABT702(+)3mix (0.1); b, *p* <0.01 vs. control; c, *p* <0.01 vs. ABT702(+); d, *p* <0.001 vs. ABT702(+)3mix(0.1); e, *p* <0.0001 vs. control; f, *p* <0.0001 vs. ABT702(+); g, *p* <0.01 vs. ABT702(+)3mix(0.1); h, *p* <0.01 vs. ABT702(+)3mix(1); i, *p* <0.05 vs. control; j, *p* <0.001 vs. ABT702; k, *p* <0.001 vs. ABT702(+)3mix(1), by one-way ANOVA followed by Tukey’s multiple comparison test. Schematic diagram showing that both lipogenesis and lipid metabolism are impaired in ADK deficiency.

